# Capacity-Contribution Paradox: Fatty Acid Oxidation Compensates for Low Glucose-Derived Acetyl-CoA in Cancer

**DOI:** 10.1101/2024.03.25.586547

**Authors:** Nancy T Santiappillai, Xiangwen C Kong, Mariam F Hakeem-Sanni, Masih Ghasemi, Anabel Withy, Lake-Ee Quek, Andrew J Hoy

**Affiliations:** School of Medical Sciences, Charles Perkins Centre, The University of Sydney, Sydney, New South Wales 2006, Australia; School of Mathematics and Statistics, Charles Perkins Centre, The University of Sydney, Sydney, New South Wales 2006, Australia

**Keywords:** Mitochondria, fatty acid oxidation, acetyl-CoA metabolism, glutamine shunt, TCA cycle, cancer metabolism

## Abstract

Upregulated fatty acid oxidation (FAO) is a hallmark of many aggressive cancers and is widely presumed to fuel the tricarboxylic acid (TCA) cycle for ATP production. However, the quantitative relationship between FAO capacity and its contribution to mitochondrial metabolism relative to other fuels remains unclear. Here, we combined parallel stable-isotope tracing with metabolic phenotyping across a diverse panel of 27 cancer cell lines to reveal a fundamental capacity-contribution paradox. Despite exhibiting FAO rates that varied over eight-fold, exogenous long-chain fatty acids consistently contributed minimally (<10%) to TCA cycle intermediates in all cancer cell lines. We demonstrated that FAO functions as a compensatory source of acetyl-CoA in cells with limited glucose-derived acetyl-CoA synthesis, rather than acting as a primary fuel source. In this context, high FAO rates do not primarily result from FAO-mediated suppression of glucose oxidation, but instead reflect the recruitment of fatty acid- and glutamine-derived carbons via a malic enzyme-dependent shunt to sustain the mitochondrial acetyl-CoA pool while preserving glucose-derived anaplerotic flux. These findings challenge the prevailing view that FAO serves as a primary bioenergetic fuel in cancer, instead identifying it as a compensatory rheostat that supplements acetyl–CoA supply in glucose–limited settings by working together with glutamine–driven, malic enzyme–dependent anaplerosis, providing a mechanistic framework to reinterpret the efficacy of FAO inhibitors beyond simple caloric starvation.

## Introduction

The integration of diverse substrates characterises cellular metabolism. Historically, glucose and glutamine have been the focal points of metabolic research in oncology, and are recognised as primary fuel sources and biosynthetic substrates ^1–3^. However, these pathways do not function in isolation; instead, they interact within an interconnected metabolic network, regulated by mitochondrial function, redox states, and nutrient sensing, among other mechanisms ^4^. Metabolic plasticity, the capacity of cancer cells to flexibly reprogram energy production and biosynthetic processes in response to changing environmental and intracellular cues, underpins both tumour survival and therapeutic resistance ^5,6^. This plasticity is intricately linked to the tricarboxylic acid (TCA) cycle, in which glucose-derived pyruvate, glutamine-derived oxoglutarate, and fatty acid-derived acetyl-CoA converge as central metabolic nodes ^4^. Understanding how these substrates compete for entry into the TCA cycle and how their utilisation is coordinated by mitochondrial biology remains profoundly underappreciated, even though this integration is central to the cancer cell phenotype and disease progression.

Fatty acid oxidation (FAO) and its interaction with mitochondrial glucose and glutamine metabolism exemplify this mechanistic gap. Genes encoding FAO enzymes are frequently upregulated in diverse cancer types, including glioma^7^, prostate^8^ and lung cancers^9^, and pharmacological inhibition of carnitine palmitoyltransferase 1 (CPT1), the rate-limiting enzyme controlling mitochondrial fatty acid entry^10^, reduces ATP production and cancer cell viability in multiple malignancies^11–18^. However, these studies presume that upregulated FAO capacity directly translates into substantial TCA cycle fuelling and bioenergetic output, an assumption rooted in classical biochemistry textbooks rather than empirical validation in cancer cells. Critically, previous investigations into cancer cell FA metabolism, including FAO, typically examined these pathways in isolation, often lacking quantitative measures of concurrent glucose or glutamine oxidation ^19–23^. Thus, they failed to account for substrate competition and metabolic plasticity, which determine whether FAO serves as a major carbon source for the TCA cycle or plays alternative metabolic roles. Collectively, these findings reveal a critical gap in our understanding of cancer metabolism: the quantitative relationship between FAO capacity and its contribution to central carbon flux relative to other major fuels, such as glucose and glutamine.

Here, we address this gap through an integrated multimodal approach combining parallel stable isotope tracing, quantitative assessment of mitochondrial biology, and metabolic correlation studies across a diverse panel of cancer cell lines spanning ten tissue origins with heterogeneous FAO capacity. Our analysis revealed a fundamental complexity in mitochondrial substrate utilisation, with exogenous long-chain fatty acids contributing minimally (<10%) to TCA cycle intermediate pools compared to glucose (30–50%) and glutamine (20–45%), despite FAO rates ranging over eight-fold among our pan-cancer cell panel. This paradox of capacity versus contribution was resolved by identifying that cells with faster FAO rates exhibit coordinated metabolic reorganisation, in which low glucose-derived acetyl-CoA synthesis is supplemented by increased recruitment of fatty acid- and glutamine-derived carbons, while glucose-derived anaplerosis is preserved. These findings suggest that FAO functions not as a static bioenergetic input but as a dynamic metabolic rheostat within an integrated network, where mitochondrial functional states tune acetyl–CoA supply by coupling fatty acid oxidation to glutamine–driven, malic enzyme–dependent acetyl–CoA production. By defining distinct functional metabolic states characterised by unique isotope-labelling signatures and mitochondrial phenotypes, we establish a quantitative framework for understanding metabolic heterogeneity as a determinant of cancer cell phenotype and therapeutic vulnerability, with implications for precision oncology and metabolic phenotyping strategies.

## Results

### Fatty acid oxidation in cancer cells showed diverse rates and spare capacity

To investigate the role of fatty acids in cancer metabolism, it is essential first to establish the baseline capacity of diverse tumour lineages to oxidise fatty acids. We first screened a panel of 27 cancer cells from 10 different origins (Figure 1A) for basal FAO rates using [1-^14^C]-palmitate and measuring CO2 production, revealing remarkable heterogeneity. In general, prostate cancer cells exhibited some of the highest FAO rates among the cancer types (Figure 1B).

**Figure 1.**
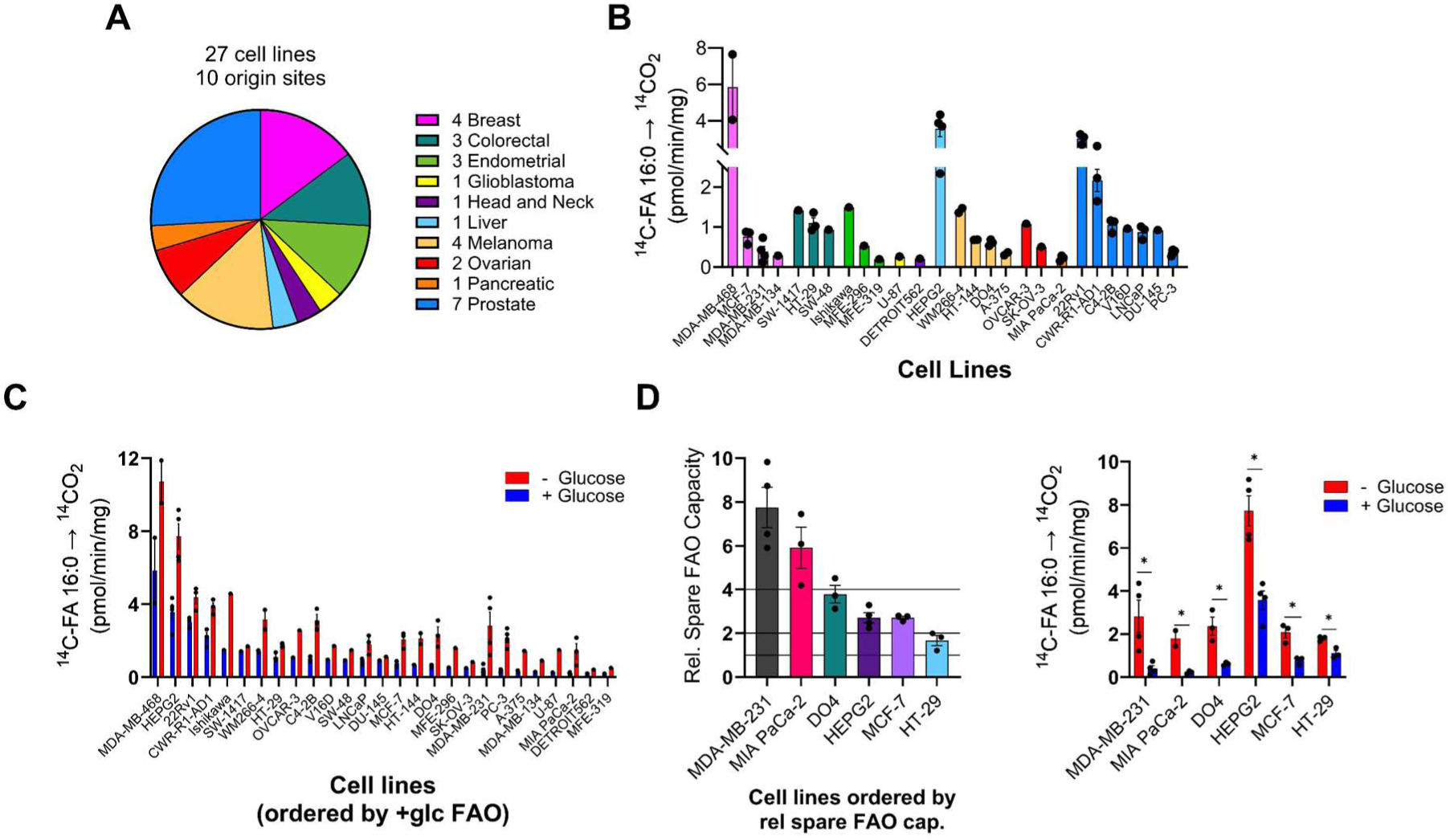
**Cancer cells exhibit heterogeneous fatty acid oxidation rates and distinct metabolic spare capacities in response to glucose availability.** (A) Panel of 27 cancer cell lines from 10 tissue origin sites measured for FAO capacity. (B) Basal [1-^14^C]-FA 16:0 oxidation rates in the presence of 5 mM glucose. Cells ordered by cancer origin site. N=1-5 per cell line. (C) [1-^14^C]-FA 16:0 oxidation rates in the presence (5 mM; blue) or absence of glucose (red). N=1-5 per cell line. (D) Selection of 6 cell lines based on relative spare FAO capacity (maximal FAO rate/basal FAO rate) differences (<2 to >7; left) between maximal FAO rate (- glucose) and basal FAO rate (+ glucose; right). N=3-4 per cell line. Graphs show mean ± SEM. * *P* < 0.05 by Student t-test. See also Figure S1.

**Figure S1, related to figure 1.**
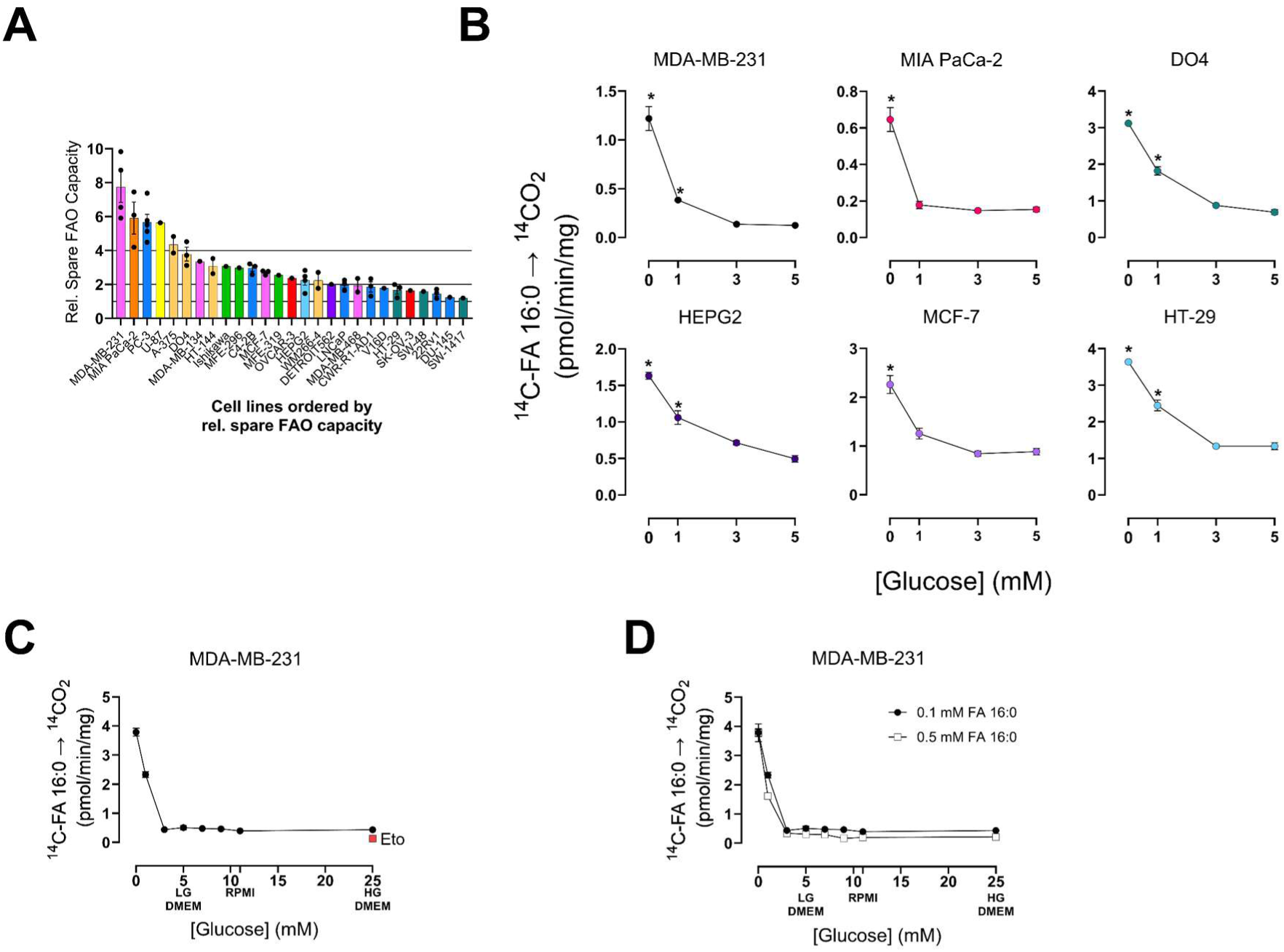
**Glucose availability suppresses fatty acid oxidation rates across diverse cancer lineages.** (A) Relative spare FAO capacity of cancer cells determined as [1-^14^C]-FA 16:0 (palmitate) oxidation rates in the absence of glucose (maximal FAO rate) divided by [1-^14^C]-FA 16:0 oxidation rates in the presence of 5 mM glucose (basal FAO rate). Cell lines ranked by fold change magnitude. N=1-5 per cell line. (B) [1-^14^C]-FA 16:0 oxidation rates in the presence of 5 mM, 3 mM, 1 mM, or 0 mM glucose. N=3 per cell line. * *P* < 0.05 vs 5 mM glucose by One-way ANOVA with Dunnett’s multiple comparisons test. (C) [1-^14^C]-FA 16:0 oxidation rates in MDA-MB-231 cells in the presence of 0 to 25 mM glucose, or 25 mM + 10 μM etomoxir, in 0.1mM ^14^C-FA 16:0. N=1 with 3 technical replicates. (D) [1-^14^C]-FA 16:0 oxidation rates in MDA-MB-231 cells in the presence of 0 to 25 mM glucose in 0.1 mM FA 16:0 compared to 0.5 mM FA 16:0. N=1 with 3 technical replicates. Graphs show mean ± SEM.

Next, we sought to gain insights into the dynamic range, or plasticity, of FAO, and thereby the “spare capacity” of this pathway to ramp up flux in response to stress. It has been shown that FAO rates increase in response to glucose metabolism inhibition in glioblastoma and breast cancer cells ^24,25^. As such, we quantified FAO rates in glucose-free conditions and termed this the maximal FAO rate. As predicted, glucose withdrawal triggered a universal upregulation of FAO across all cancer lines tested (Figure 1C and S1A), with increases ranging from 2-fold to > 8-fold. Based on the fold change in maximal and basal FAO rates as a measure of relative FAO spare capacity, we selected six cell lines that ranged from high (> 4-fold), mid (2-4-fold), and low (0-2-fold) FAO spare capacity: MDA-MB-231 and MCF-7 (breast), MIA PaCa-2 (pancreatic), DO4 (melanoma), HEPG2 (liver), and HT-29 (colorectal) (Figure 1D). We further explored the relationship between FAO rates and media glucose levels in this panel of cells and determined that FAO is actively suppressed by glucose levels of 3 mM or greater (Figure S1B and S1C). Conversely, FAO was not dependent on nutrient gradients since a 5-fold increase in media palmitate (0.1 mM to 0.5 mM) did not increase FAO rate in the presence of 5 mM glucose (Figure S1D). These data are consistent with a model in which FAO is preferentially engaged when glucose utilisation is limited, either by availability or intrinsic constraints on glucose-derived acetyl–CoA synthesis.

### Palmitate is a minor TCA cycle carbon source compared to glucose and glutamine

Next, we used a parallel tracing strategy that fed cells uniformly labelled with ^13^C-glucose, glutamine, or palmitate (FA 16:0) to gain new insights into the quantitative contributions of enrichment to TCA cycle intermediates in our subset of six cell lines spanning FAO spare capacity (Figure 2A). We first identified that 6 hours was the optimal tracing duration, as media substrate levels were not fully depleted and there was no evidence of impaired substrate uptake (Figure S2A). This was critical because we showed that removal or very low levels of extracellular glucose had a striking effect on FAO rates (Figure 1). As such, the depletion of glucose from the media after 12 and 24 hours likely affects ^13^C-palmitate metabolism and enrichment patterns. This rapid rate of media glucose consumption can somewhat be avoided by doubling the media volume (Figure S2B).

**Figure 2.**
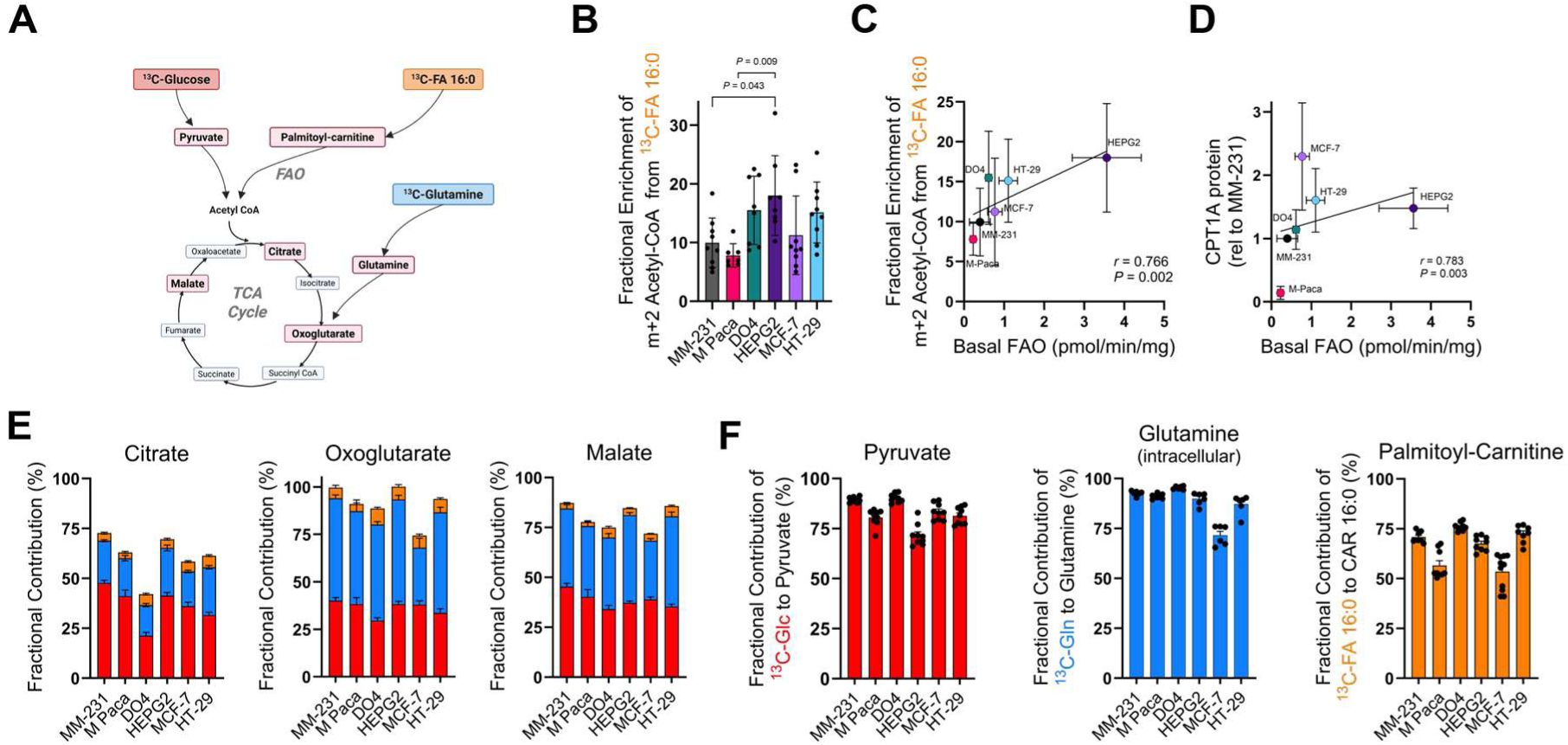
**Exogenous fatty acids are minor carbon contributors to the TCA cycle compared to glucose and glutamine, irrespective of oxidative capacity.** (A) Schematic representation of [U-^13^C]-glucose (5 mM), glutamine (1 mM), or FA 16:0 (150 μM) tracing into the TCA cycle and precursor metabolite. (B) Fractional contribution of [U-^13^C]-FA 16:0 to m+2 acetyl-CoA after 6 hours of labelling. N=9 per cell line. (C) Correlation between [U-^13^C]-FA 16:0 to m+2 acetyl-CoA and basal FAO rate. Error bars represent ± SD. *P* and *r* values by linear regression. (D) Correlation between CPT1a protein levels and basal FAO rate. Error bars represent ± SD. *P* and *r* values by linear regression. (E) Fractional contributions of [U-^13^C]-glucose (red), glutamine (blue), or FA 16:0 (orange) to citrate, oxoglutarate, and malate. N=9 per cell line.(F) Fractional contribution of [U-^13^C]-glucose to pyruvate (red), [U-^13^C]-glutamine to intracellular glutamine (blue), [U-^13^C]-FA 16:0 to palmitoyl-carnitine (orange) after 6 hours of labelling. N=9 per cell line. Level of natural enrichment of metabolite standards indicated. Graphs show mean ± SEM unless stated otherwise. ^13^C-FA 16:0: U-^13^C-palmitate; MM-231: MDA-MB-231; M-Paca: Mia Paca-2. See also Figure S2.

**Figure S2, related to figure 2.**
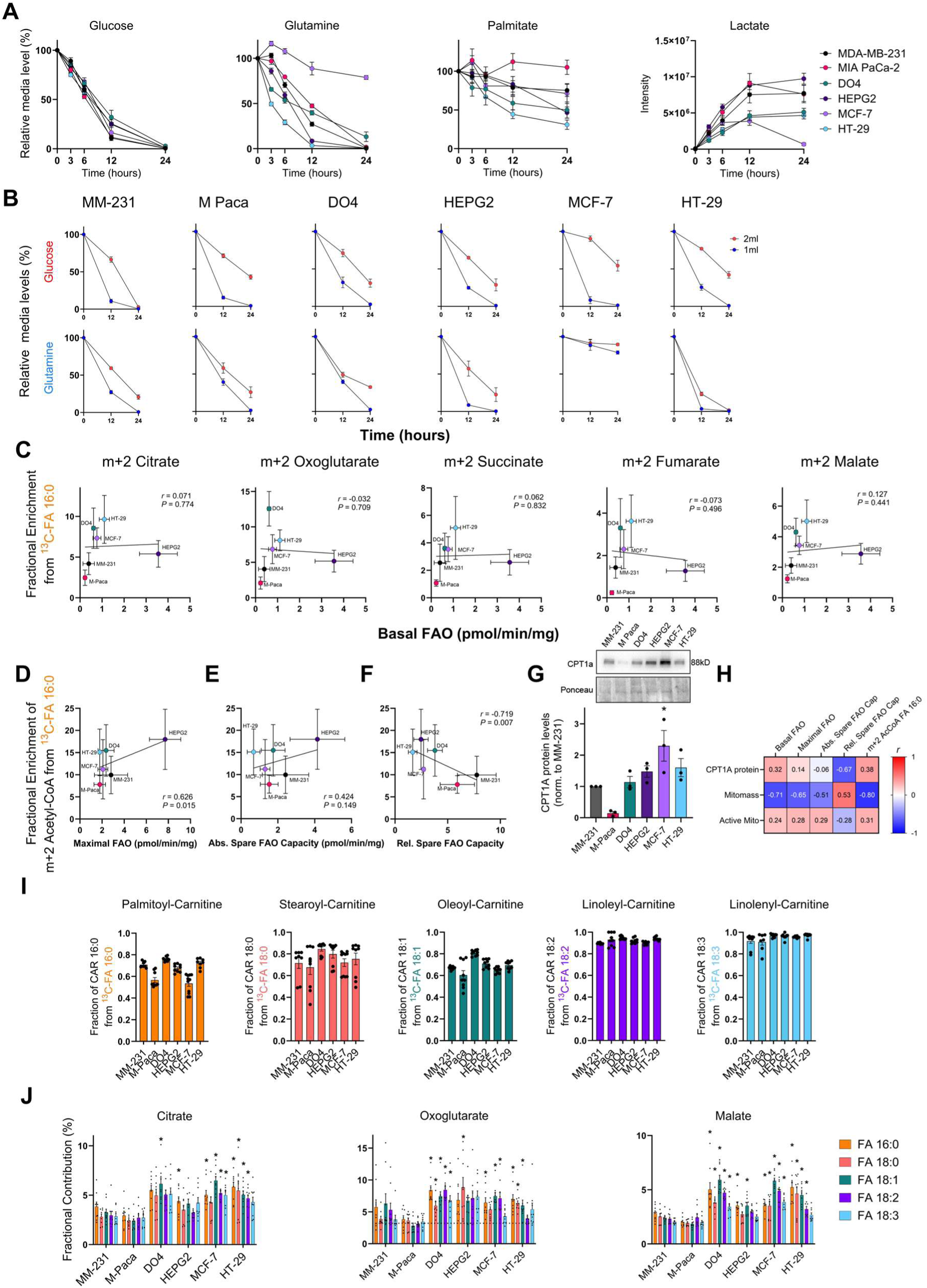
**Kinetic profiling of substrate uptake and lack of correlation between mitochondrial mass and FAO capacity.** (A) Media glucose, glutamine, FA 16:0 (palmitate) levels relative to t = 0, and lactate intensity over 24 hours. N=3 per cell line. (B) Media glucose or glutamine levels over 24 hours relative to t = 0 from wells containing 1 mL (blue) or 2 mL (red) of medium. N=3 per cell line. Graphs show mean ± SEM (C) Fractional contribution of [U-^13^C]-FA 16:0 to m+2 TCA cycle metabolites after 6 hours of labelling. N=9 per cell line. Level of natural enrichment of metabolite standards indicated. Graphs show mean ± SD. *P* and *r* values by linear regression. (D) Correlation between [U-^13^C]-FA 16:0 to m+2 acetyl-CoA and maximal FAO rate (glucose-free media). Error bars represent ± SD. *P* and *r* values by linear regression. (E) Correlation between [U-^13^C]-FA 16:0 to m+2 acetyl-CoA and absolute spare FAO rate (Maximal FAO – Basal FAO). Error bars represent ± SD. *P* and *r* values by linear regression. (F) Correlation between [U-^13^C]-FA 16:0 to m+2 acetyl-CoA and relative spare FAO rate (Maximal FAO / Basal FAO). Error bars represent ± SD. *P* and *r* values by linear regression. (G) Representative immunoblots and densitometric quantitation of CPT1a. (H) Correlation matrix of measures of FAO capacity and CPT1 protein level, and mitochondrial mass (using MitoTracker Green) and mitochondrial membrane potential (Active Mito; using MitoTracker Red). *r* values by linear regression. (I) Fractional contributions of [U-^13^C]- FA 16:0 (palmitate), 18:0 (stearate), 18:1 (oleate), 18:2 (linoleate), and 18:3 (linolenate) to acyl-carnitine species. N=9 per cell line. (J) Fractional contributions of [U-^13^C]- FA 16:0 (palmitate), 18:0 (stearate), 18:1 (oleate), 18:2 (linoleate), and 18:3 (linolenate) to citrate, oxoglutarate, and malate. N=9 per cell line. Graphs show mean ± SEM unless stated otherwise. ^13^C-FA 16:0: U-^13^C-palmitate; MM-231: MDA-MB-231; M-Paca: Mia Paca-2.

We used m+2 acetyl-CoA from U-^13^C-palmitate as our stable-isotope measure of FAO (Figure 2B), which, as expected, positively correlated with basal FAO rate (*r* = 0.77; Figure 2C), as determined using ^14^C-palmitate tracing (Figure 1). However, basal FAO rate did not correlate with ^13^C-palmitate enrichment of other TCA cycle intermediates (Figure S2C). The fractional contribution of ^13^C-palmitate to m+2 acetyl-CoA did positively correlate with maximal FAO rate (glucose-free media; Figure S2D), not with the absolute FAO spare capacity (delta between maximal and basal FAO rate; Figure S2E), but interestingly, there was a negative correlation with the relative FAO spare capacity (maximal FAO/basal FAO; Figure S2F). These observations suggest that cells with high basal FAO (and high m+2 acetyl-CoA from palmitate) operate closer to their mitochondrial FAO ceiling and exhibit limited relative plasticity for FAO upregulation. In contrast, low-basal FAO cells retain greater proportional reserve capacity despite lower absolute rates. Critically, these data highlight that basal FAO rate does not predict the FAO rate or capacity in other settings and that there is significant heterogeneity in both relative and absolute FAO spare capacity in cancer cells.

These patterns of fatty acid metabolism in our panel of cancer cells were, in part, explained by features of mitochondrial biology. Cells with higher basal FAO rates had greater CPT1A protein levels (Figure 2D and S2G), but this did not predict maximal FAO capacity or the relative spare capacity of the cells, as there was no relationship between CPT1A protein levels and maximal FAO capacity or relative spare capacity (Figure S2H). In fact, cells with greater relative spare FAO capacity had lower CPT1 protein levels (*r* = −0.665, *P* = 0.013; Figure 2H). This distinction is vital for interpreting therapeutic targets; inhibition of CPT1A may have disproportionate effects depending on whether a tumour is operating at basal or maximal capacity. Furthermore, cells with high FAO rates frequently exhibited lower mitochondrial mass (quantified using MitoTracker Green; Figure S2H). Finally, there was no relationship between measures of FAO and mitochondrial membrane potential, determined using MitoTracker Red (normalised to MitoTracker Green; Figure S2H). These data imply that high basal FAO flux is achieved through higher CPT1 protein levels and likely greater efficiency, rather than through global mitochondrial biogenesis.

A notable outcome of the U-^13^C-palmitate tracing experiments was the very low enrichment of acetyl-CoA and TCA cycle metabolites, with palmitate minimally contributing 9-18% of carbons to acetyl-CoA (Figure 2B), and <6% to TCA cycle metabolites citrate, oxoglutarate and malate, compared to glucose (30-50%) and glutamine (20-45%) (Figure 2E). This is, even though FAO is commonly assumed to be a major contributor to the TCA cycle by producing acetyl-CoA and reducing equivalents to support ATP production ^26,27^. This low rate of enrichment of acetyl-CoA, citrate and other TCA cycle intermediates was not due to low precursor enrichment (>56% enrichment of palmitoyl-carnitine from ^13^C-palmitate) (Figure 2F), nor did differences in precursor labelling explain differences in the contribution of glucose and glutamine; >80% enrichment of pyruvate from ^13^C-glucose, and >60% enrichment of intracellular glutamine from extracellular ^13^C-glutamine (Figure 2F). Furthermore, this low enrichment was also observed with other FAs that varied in saturation (double bonds) and carbon chain lengths; specifically [U-^13^C]-stearate (FA 18:0), oleate (FA 18:1), linoleate (FA 18:2), and linolenate (FA 18:3). In all experiments, each ^13^C-FA enriched their direct acyl-carnitine counterpart species by a fractional ratio of 0.5-0.95 (Figure S2I), and exogenous LCFAs contributed <10% of labelled carbons to citrate, oxoglutarate, or malate (Figure S2J). Collectively, these data further demonstrate that exogenous long-chain fatty acids, regardless of saturation status or chain length, are minor carbon sources for the TCA cycle across a pan-cancer panel of cells spanning a range of FAO rates and spare capacities. As such, these findings directly challenge the fuel hypothesis of the role of FAO in cancer cells, because if FAO accounts for a significant portion of oxygen consumption but provides <10% of the TCA cycle carbon, the pathway cannot serve as the primary source of biomass or bioenergetic carbon.

### Fatty acid oxidation compensates for reduced glucose-derived acetyl-CoA in the mitochondrial pool

Given the extensive interest in FAO in cancer ^28–31^, and our observations that FAO is not a major carbon source for fuelling the TCA cycle, we sought to gain new insights into how cancer cells integrate mitochondrial metabolic activity in our panel of cancer cell lines with varying FAO capacity. We first focused on glucose metabolism, given the seminal work of Randle and colleagues ^32^, which showed the interaction between fatty acid and glucose metabolism ^32^, their shared input into the mitochondrial acetyl-CoA pool, and the relationship between glucose availability and FAO rates (Figure 1 and S1). Cells with high basal and maximal FAO rates had higher glucose exclusion, as indicated by m+0 (Figure 3A), and lower m+2 acetyl-CoA (Figure 3B) labelling from [U-^13^C]-glucose, indicating reduced glucose-derived acetyl-CoA synthesis. This relationship between FAO rate and glucose contribution to acetyl-CoA was supported by very strong negative correlations between the contribution of palmitate and glucose to m+2 acetyl-CoA (*r* = −0.98; Figure 3C), which was also observed in cells cultured with [U-^13^C]-stearate (*r* = −0.95; Figure 3D), oleate (*r* = −0.76; Figure 3E), linoleate (*r* = −0.89; Figure 3F), or linolenate (*r* = −0.97; Figure 3G). These data indicate that in cells where glucose contributes less to acetyl-CoA, fatty acid-derived carbons proportionally contribute more, consistent with a compensatory relationship between these substrates at the level of the acetyl-CoA pool. This inverse relationship between the contribution of glucose and fatty acid to acetyl-CoA was also evident in citrate (m+2; *r* = −0.43; Figure 3H), but not in m+2 labelling of oxoglutarate (Figure 3I), succinate (Figure 3J), fumarate (Figure 3K) or malate (Figure 3L). Overall, we observe a Randle Cycle-like relationship between glucose and fatty acid metabolism at acetyl-CoA in our panel of cancer cells, but this was not propagated through TCA cycle intermediates downstream of glutamine entry. Further, these data support a model in which low glucose-derived acetyl–CoA is compensated by FAO, rather than FAO being the primary driver of glucose suppression

**Figure 3.**
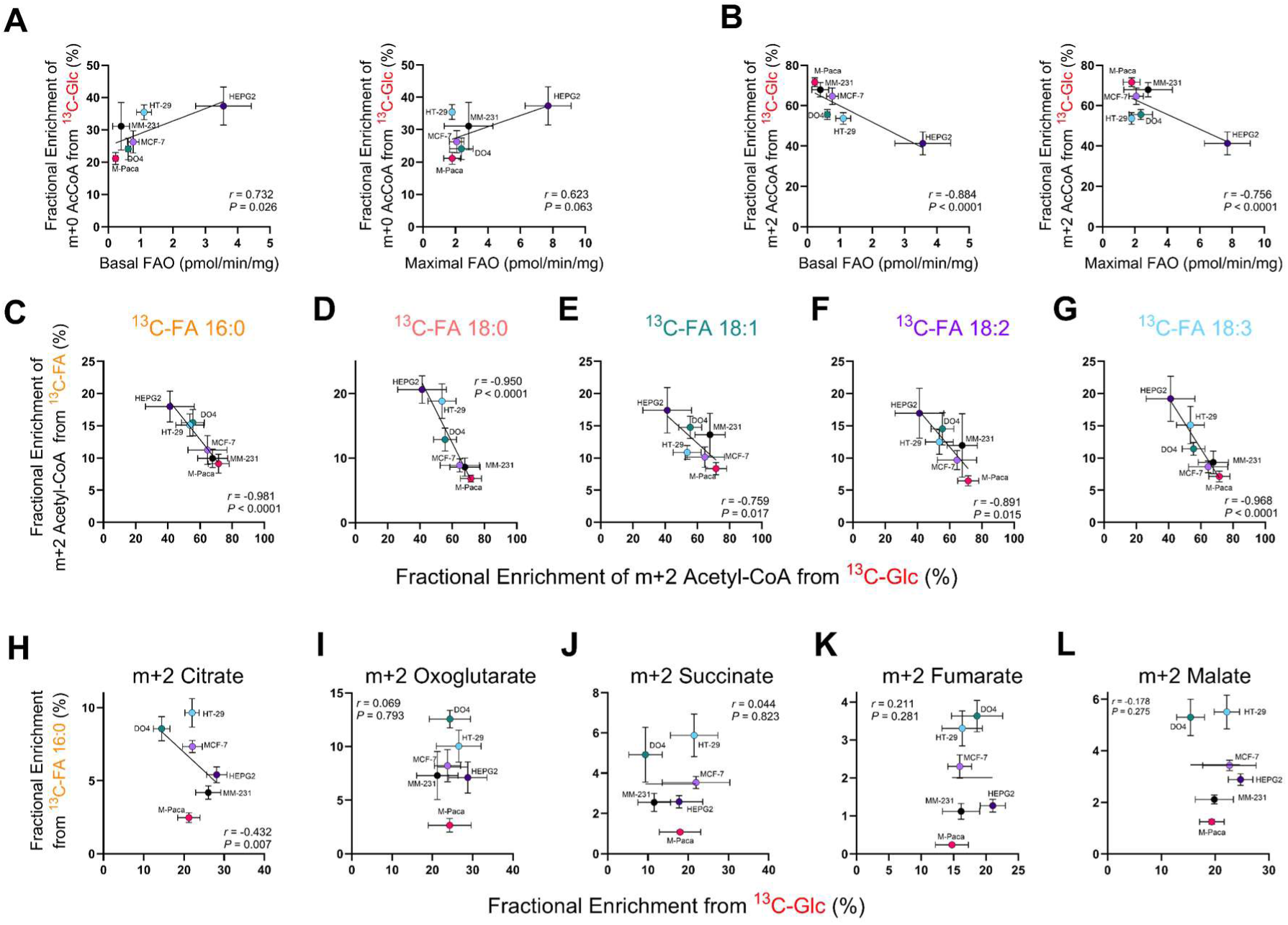
**Fatty acid oxidation compensates for reduced glucose-derived acetyl-CoA, consistent with the Randle Cycle.** (A) Correlation between m+0 acetyl-CoA in cells cultured in 5 mM [U-^13^C]-glucose and basal (left) and maximal (right) FAO rates. (B) Correlation between m+2 acetyl-CoA in cells cultured in 5 mM [U-^13^C]-glucose and basal (left) and maximal (right) FAO rates. (C) Correlation between [U-^13^C]-FA 16:0 (palmitate) and [U-^13^C]-glucose contributions to m+2 acetyl-CoA. (D) Correlation between [U-^13^C]-FA 18:0 (stearate) and [U-^13^C]-glucose contributions to m+2 acetyl-CoA. (E) Correlation between [U-^13^C]-FA 18:1 (oleate) and [U-^13^C]-glucose contributions to m+2 acetyl-CoA. (F) Correlation between [U-^13^C]-FA 18:2 (linoleate) and [U-^13^C]-glucose contributions to m+2 acetyl-CoA. (G) Correlation between [U-^13^C]-FA 18:3 (linolenate) and [U-^13^C]-glucose contributions to m+2 acetyl-CoA. (H) Correlation between [U-^13^C]-FA 16:0 and [U-^13^C]-glucose contributions to m+2 citrate. (I) Correlation between [U-^13^C]-FA 16:0 and [U-^13^C]-glucose contributions to m+2 oxoglutarate. (J) Correlation between [U-^13^C]-FA 16:0 and [U-^13^C]-glucose contributions to m+2 succinate. (K) Correlation between [U-^13^C]-FA 16:0 and [U-^13^C]-glucose contributions to m+2 fumarate. (L) Correlation between [U-^13^C]-FA 16:0 and [U-^13^C]-glucose contributions to m+2 malate. Graphs show mean ± SD. *P* and *r* values by linear regression. ^13^C-FA 16:0: U-^13^C-palmitate; ^13^C-Glc: U-^13^C-glucose; MM-231: MDA-MB-231; M-Paca: Mia Paca-2.

### A compensatory ’dual-fuel’ shunt recruits glutamine to maintain acetyl-CoA availability in high-FAO phenotypes

Having established that higher FAO rates are associated with reduced glucose-derived acetyl–CoA labelling (consistent with a Randle cycle-like relationship at the acetyl–CoA node), which does not extend into TCA cycle intermediates, we next investigated the relationship between FAO and glutamine metabolism (Figure 4A). In striking contrast to the glucose data, cancer cells with high FAO rates exhibited significantly higher glutamine oxidation rates, as evidenced by a strong positive correlation with m+4 succinate from ^13^C-glutamine (*r* = 0.90; Figure 4B). This positive coupling was consistent across all metrics of FAO potential, including maximal FAO capacity (*r* = 0.88; Figure 4C), and absolute FAO spare capacity (e.g. maximal FAO – basal FAO, *r* = 0.75; Figure S4A), but not relative spare FAO capacity (e.g. maximal FAO/basal FAO ratio; Figure S4B). There was also a coupling of ^13^C-palmitate contribution to m+2 acetyl-CoA with m+4 succinate from ^13^C-glutamine (*r* = 0.49; Figure 4D). This pattern extended throughout the TCA cycle and downstream amino acids, with significant positive correlations observed between FAO activity and glutamine labelling in fumarate, malate, and aspartate (Figure S4C). These data suggest that, rather than competing for mitochondrial entry, FAO and glutamine oxidation are coordinately upregulated and, as such, predict that high FAO cells exhibit low glutamine reductive carboxylation. Surprisingly, cells with higher FAO rates showed increased glutamine reductive carboxylation (indicated by m+5 citrate (Figure 4E) and m+3 aspartate (Figure S4D)). However, when normalising for oxidative flux, cells with higher FAO rates displayed a reduced relative reliance on reductive carboxylation (lower m+5/m+4 citrate ratio; *r* = −0.336, *P* = 0.024; Figure 4F). Therefore, cells with higher FAO and lower glucose oxidation preferentially maintain canonical oxidative glutamine cycling. Notably, these cells did not exhibit altered glucose anaplerosis compared with low FAO cells, and there was no correlation between FAO rates and glucose entry via pyruvate carboxylase (m+3 citrate from ^13^C-glc; Figure S4E) or between FAO rates and pyruvate partitioning between anaplerosis and cataplerosis (m+3/m+2 citrate from ^13^C-glc; Figure S4F).

**Figure 4.**
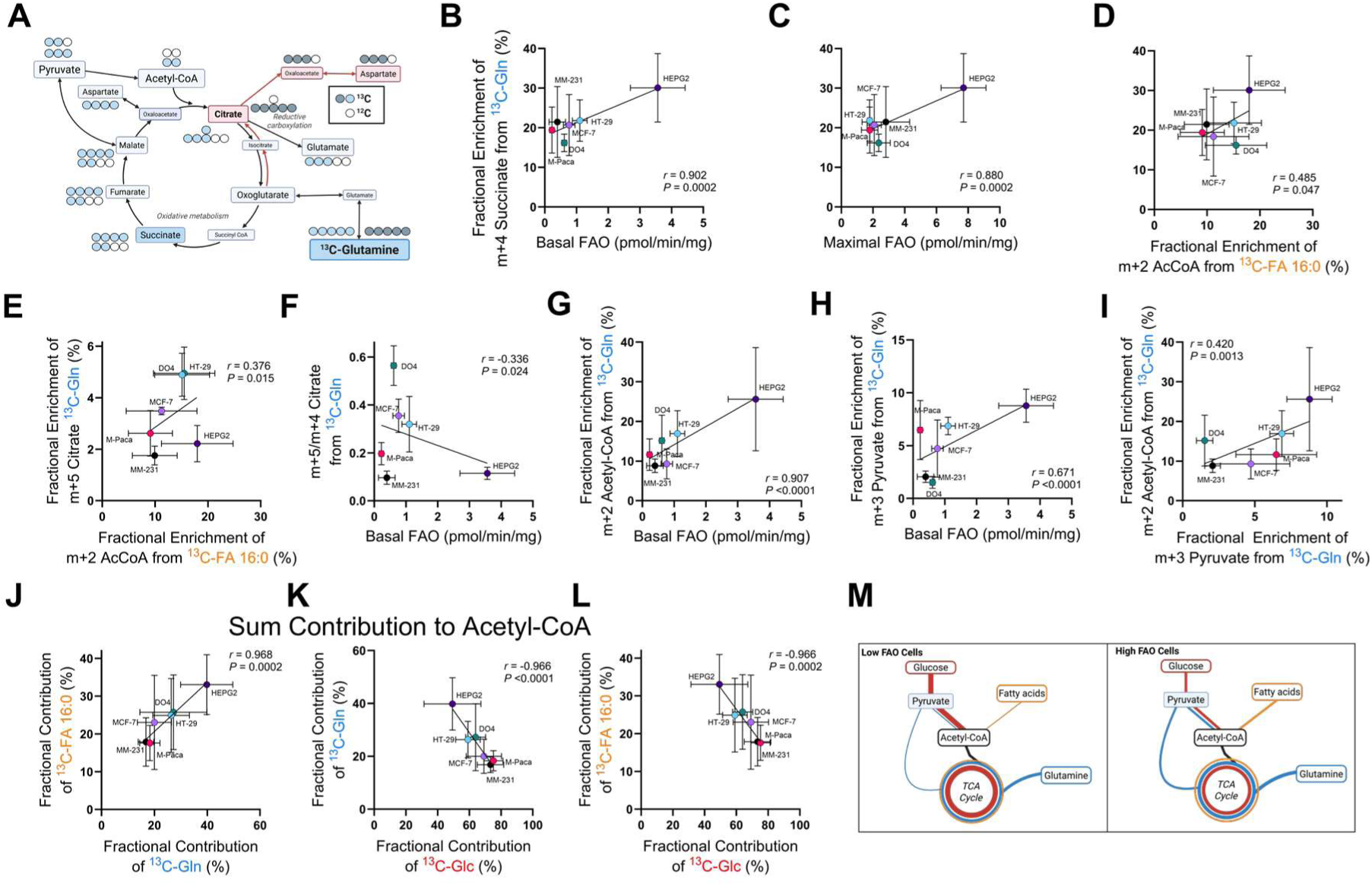
**High fatty acid oxidation rates are positively coupled to glutamine metabolism and drive a compensatory dual-fuel shunt to maintain acetyl-CoA.** (A) Schematic of [U-^13^C]-glutamine (^13^C-Gln) oxidation and reductive carboxylation. (B) Correlation between m+4 succinate from glutamine (glutamine oxidation) and basal FAO rate. (C) Correlation between m+4 succinate from glutamine (glutamine oxidation) and maximal FAO rate. (D) Correlation between m+4 succinate from glutamine (glutamine oxidation) and m+2 acetyl-CoA from [U-^13^C]-FA 16:0 (palmitate). (E) Correlation between m+5 citrate from glutamine (reductive carboxylation) and m+2 acetyl-CoA from [U-^13^C]-FA 16:0 (palmitate). (F) Correlation between m+5 citrate / m+4 citrate from glutamine (reductive carboxylation / glutamine oxidation) and basal FAO rate. (G) Correlation between m+3 pyruvate from glutamine and basal FAO rate. (H) Correlation between m+2 acetyl-CoA from glutamine and basal FAO rate. (I) Correlation between m+3 pyruvate and m+2 acetyl-CoA from glutamine. (J) Correlation between [U-^13^C]-FA 16:0 and [U-^13^C]-glutamine to acetyl-CoA. (K) Correlation between [U-^13^C]-glutamine and [U-^13^C]-glucose (^13^C-Glc) to acetyl-CoA. (L) Correlation between [U-^13^C]-FA 16:0 and [U-^13^C]-glucose to acetyl-CoA. (M) Schematic of dual-fuel link between FAO and glutamine metabolism to maintain acetyl-CoA levels. Graphs show mean ± SD. *P* and *r* values by linear regression. ^13^C-FA 16:0: U-^13^C-palmitate; ^13^C-Glc: U-^13^C-glucose; ^13^C-Gln: U-^13^C-glutamine; MM-231: MDA-MB-231; M-Paca: Mia Paca-2. See also Figure S4.

**Figure S4, related to figure 4.**
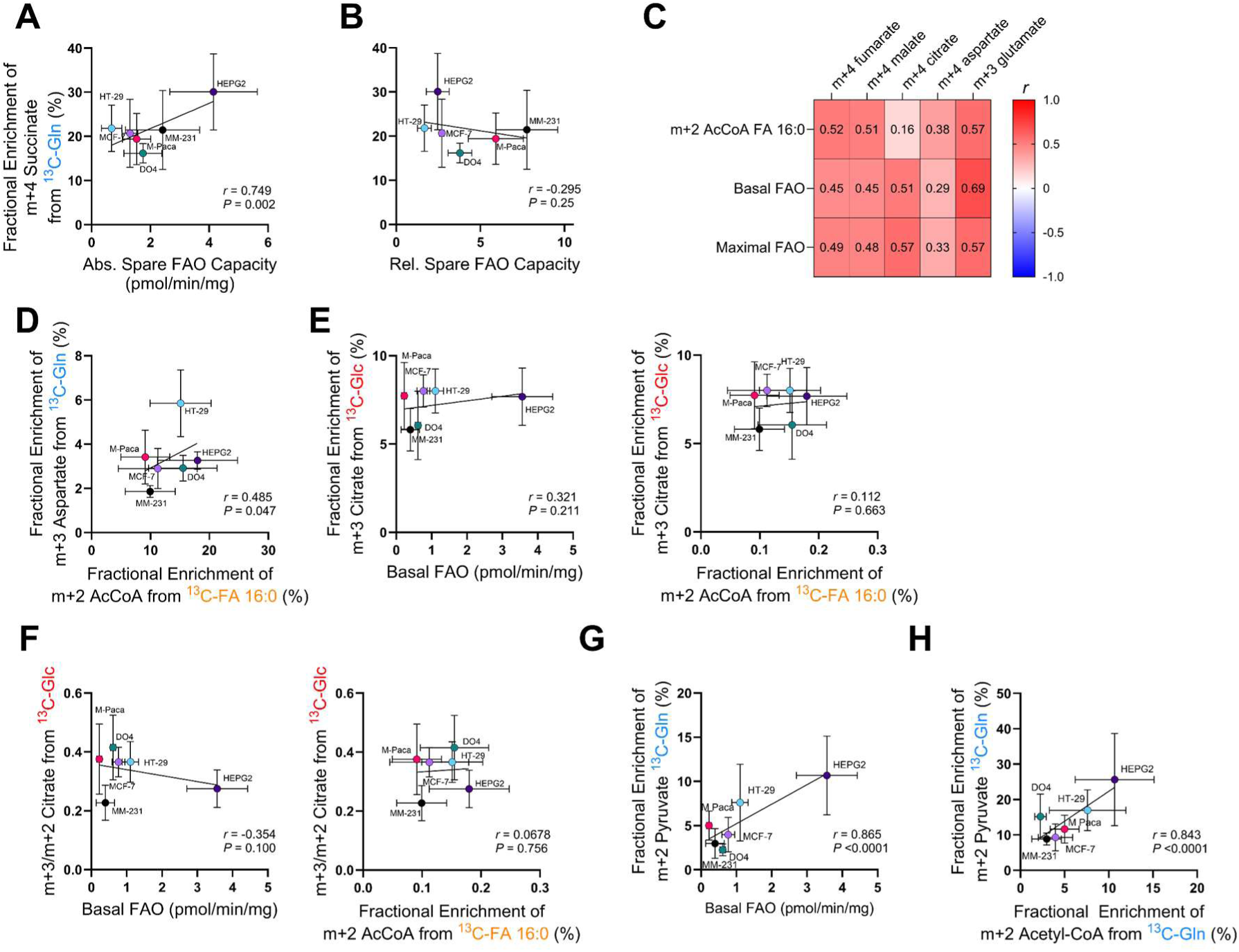
**Relationships between fatty acid oxidation capacity, glutamine metabolism and glucose anaplerotic flux.** (A) Correlation between m+4 succinate from glutamine (glutamine oxidation) and absolute spare FAO rate (Maximal FAO – Basal FAO). (B) Correlation between m+4 succinate from glutamine and relative spare FAO rate (Maximal FAO / Basal FAO). (C) Correlation matrix between [U-^13^C]-glutamine enrichment into m+4 isotopologues of metabolites and measures of FAO. (D) Correlation between m+3 aspartate from glutamine and m+2 acetyl-CoA from [U-^13^C]-FA 16:0 (palmitate). (E) Correlation between m+3 citrate from glucose (anaplerosis) and basal FAO rate (left) and [U-^13^C]-FA 16:0 to m+2 acetyl-CoA (right). (F) Correlation between m+3 citrate / m+2 citrate from glucose (anaplerosis / oxidation) and basal FAO rate (left) and [U-^13^C]-FA 16:0 to m+2 acetyl-CoA (right). (G) Correlation between m+2 pyruvate from glutamine (2^nd^ turn of TCA cycle of glutamine oxidation) and basal FAO rate. (H) Correlation between m+2 pyruvate and m+2 acetyl-CoA from glutamine. Graphs show mean ± SD unless stated otherwise. *P* and *r* values by linear regression. ^13^C-FA 16:0: U-^13^C-palmitate; ^13^C-Glc: U-^13^C-glucose; ^13^C-Gln: U-^13^C-glutamine; MM-231: MDA-MB-231; M-Paca: Mia Paca-2.

What we could not reconcile from these relationships was the absolute carbon contributions to key metabolites, especially to the synthesis of acetyl-CoA and citrate (Figure 2). For example, palmitate contributes only 9-18% of carbon to acetyl-CoA (Figure 2B) and <6% to citrate (Figure 2F) in our panel of cells, whereas glucose contributes 41-72% to acetyl-CoA (Figure 3) and 30-50% to citrate (Figure 2F). Notably, in cells with high FAO rates, fatty acids and glucose together accounted for only ∼50% of acetyl-CoA labelling, compared with ∼80% in low-FAO cells, implying that an additional substrate must sustain acetyl-CoA supply in the high-FAO state. The coupling of FAO and glutamine oxidation and reductive carboxylation Figure 4B-F) raised the prospect that this could extend the acetyl-CoA pool, where glutamine carbons enter the TCA cycle, exit via malic enzyme as pyruvate, and re-enter as acetyl–CoA, effectively backfilling the acetyl–CoA pool when glucose-derived acetyl–CoA is low in high-FAO cells. This concept was supported by a strong positive correlation between basal FAO rates and the contribution of glutamine to acetyl-CoA (m+2; r = 0.91; Figure 4G), which ranged from 9% to 26%. We saw the same positive relationship between FAO rate and glutamine labelling of pyruvate (m+3, *r* = 0.67; Figure 4H; m+2, *r* = 0.86; Figure S4G). The enrichment of glutamine-derived pyruvate was predictive of glutamine-derived acetyl-CoA (Figure 4I and S4H). More broadly, this increased enrichment of acetyl-CoA from glutamine in cells with higher FAO was also evident when we quantified the total fractional contribution from glutamine and palmitate (*r* = 0.968; Figure 4J), whereas there was a negative relationship between glucose enrichment of acetyl-CoA and glutamine (*r* = −0.966; Figure 4K) and palmitate (*r* = −0.966; Figure 4L). Collectively, these data reveal a dual-fuel mechanism in high-FAO cancer cells in which fatty acid oxidation and a malic enzyme-dependent glutamine–pyruvate shunt jointly supplement acetyl-CoA production when glucose-derived acetyl-CoA synthesis is low, while glucose-derived anaplerosis is maintained (Figure 4M).

### High FAO capacity does not confer bioenergetic autonomy under nutrient stress

Having established that FAO is a minor direct carbon source for the TCA cycle and instead supplements acetyl-CoA production alongside glutamine when glucose-derived acetyl-CoA synthesis is low, we next asked whether this compensatory role is fixed or conditional. Specifically, we questioned whether the low fractional contribution of fatty acids (<10%) in nutrient-replete conditions reflects an actively maintained metabolic configuration that can be overridden when glucose and glutamine are scarce, or whether it reveals a fundamental limitation of the cancer mitochondrial architecture. If FAO truly operates as a reserve fuel source, then deprivation of glucose and glutamine would be expected to force a switch to fatty acid dependence, converting the high FAO enzymatic capacity observed in our panel (Figure 1) into substantial TCA cycle fuelling. To test this, we cultured cells in glucose-free or glucose/glutamine-free conditions. As expected, glucose withdrawal triggered a compensatory upregulation in FAO rates (Figure 1C and 1D) and increased the fractional enrichment of ^13^C-palmitate into citrate (Figure 5A and 5B), as well as oxoglutarate and malate (Figure S5A). However, this increase was modest; even under extreme nutrient stress, the contribution of palmitate carbons to the TCA cycle rarely exceeded 15%. This increased fuelling of the TCA cycle by FAO in glucose-/glutamine-free conditions was also associated with other metabolic changes. Specifically, we observed an increase in m+2 and, to a greater extent, m+4 labelling of citrate, corresponding to the 1^st^ and 2^nd^ turns of the TCA cycle, respectively (Figure 3C). These results suggest that palmitate-derived carbons undergo multiple rounds of the TCA cycle when glucose or both glucose and glutamine are absent from the culture medium.

**Figure 5.**
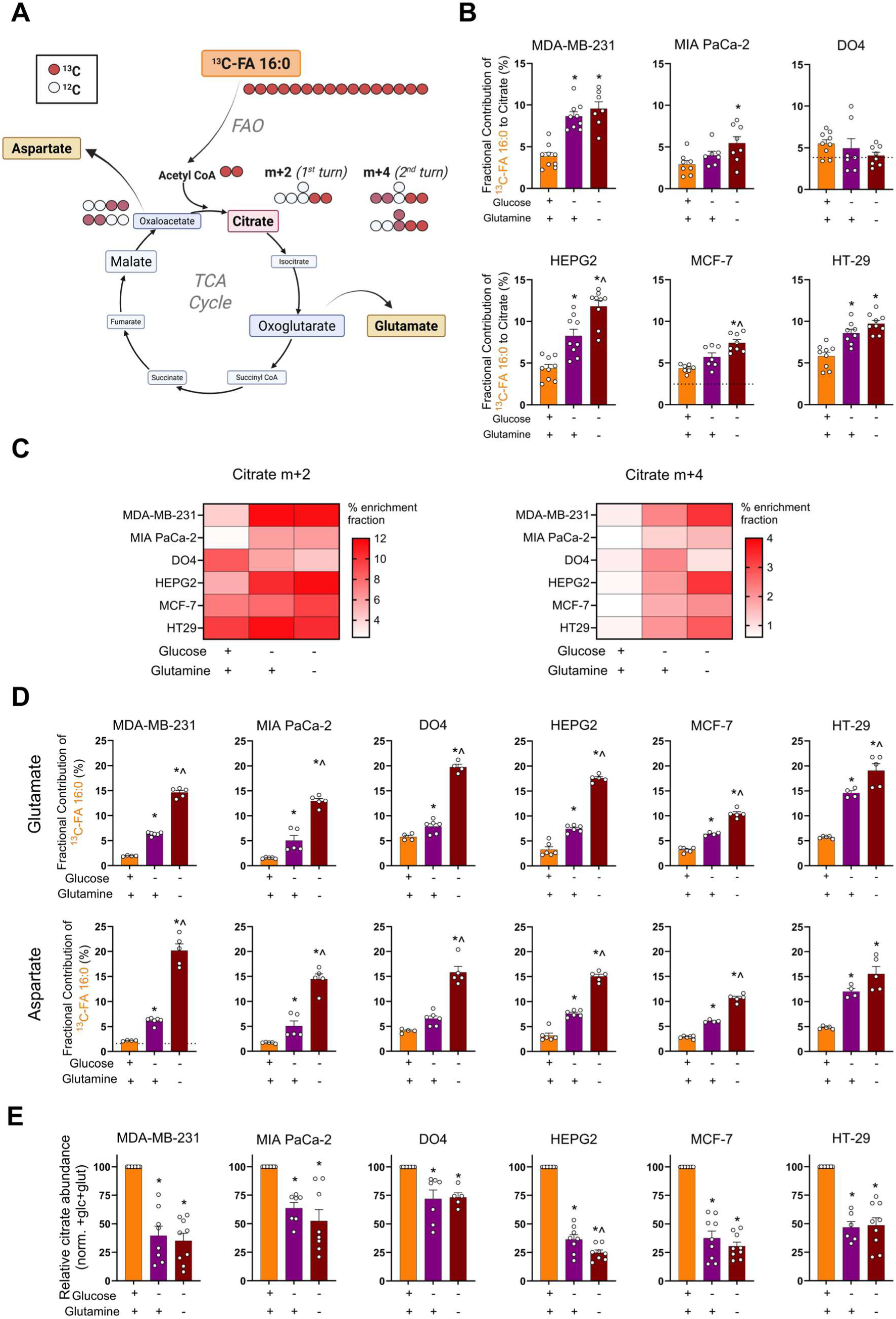
**Fatty acid oxidation is insufficient to rescue TCA cycle pool sizes or cell viability in the absence of glucose and glutamine anaplerosis.** (A) Schematic tracing the conversion of [U-^13^C]-FA 16:0 (palmitate) to citrate m+2 and m+4 glutamate, and aspartate. (B) Fractional contribution of [U-^13^C]-FA 16:0 to citrate in cells cultured in +glucose/+glutamine, - glucose/+glutamine, or -glucose/-glutamine. N=9 per cell line. (C) Enrichment fraction (%) of [U-^13^C]-FA 16:0 to m+2 and m+4 citrate in cells cultured in + glucose/+glutamine, -glucose/+glutamine, and -glucose/-glutamine. N=9 per cell line. (D) Fractional contribution of [U-^13^C]-FA 16:0 to glutamate and aspartate in cells cultured in +glucose/+glutamine, -glucose/+glutamine, or -glucose/-glutamine. N=4-6 per cell line. (E) Relative abundance of citrate in cells cultured in +glucose/+glutamine, -glucose/+glutamine, or -glucose/-glutamine, normalised to +glucose/+glutamine. N=6-9 per cell line. Graphs show mean ± SEM. * *P* < 0.05 compared to +glucose/+glutamine, ^ *P* < 0.05 compared to - glucose/+glutamine by One-way ANOVA with Dunnett’s multiple comparisons test. See also Figure S5.

**Figure S5, related to figure 5.**
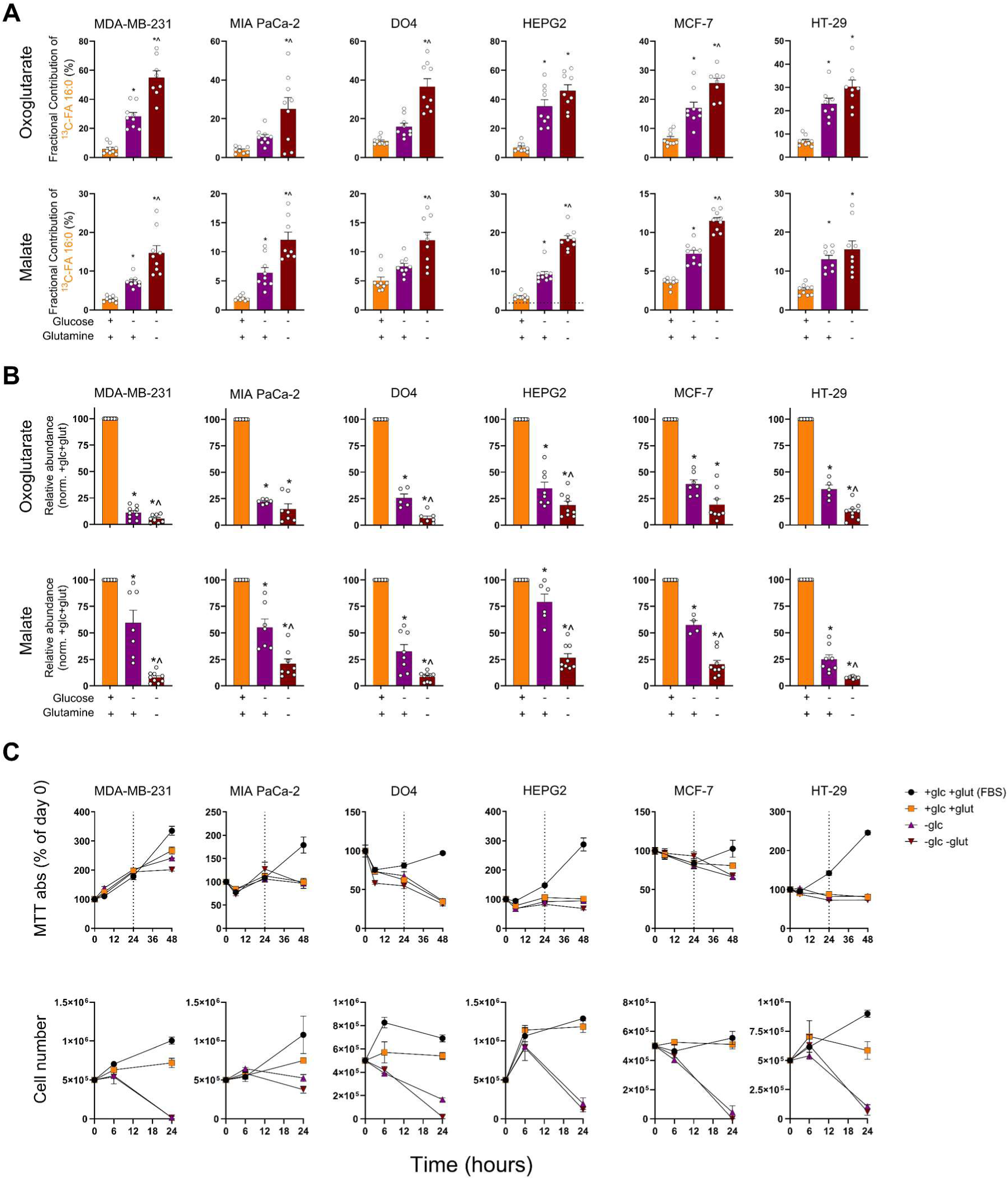
**TCA cycle intermediate abundance and cell viability under nutrient-deprived conditions.** (A) Fractional contribution of [U-^13^C]-FA 16:0 to oxoglutarate and malate in cells cultured in +glucose/+glutamine, -glucose/+glutamine, or -glucose/-glutamine. N=4-6 per cell line. (B) Relative abundance (normalised to +glucose/+glutamine) of [U-^13^C]-FA 16:0 to oxoglutarate or malate. N=5-9 per cell line. (C) Impact of glucose- or glucose and glutamine- deprivation on metabolic activity (top) assessed by MTT assay. Absorbance results up to 48 hours normalised to day 0 (dotted line at 24 hours indicates replenishment of media); or cell viability (bottom) assessed by live cell counts at 0, 6 and 24 hours. N=3 per group. Graphs show mean ± SEM. * *P* < 0.05 compared to +glucose/+glutamine, ^ *P* < 0.05 compared to - glucose/+glutamine by One-way ANOVA with Dunnett’s multiple comparisons test. Glc: glucose; gln: glutamine.

Another striking shift in palmitate metabolism in glucose-/ glutamine-deprived conditions was the enrichment of the cataplerotic metabolites, glutamate and aspartate (Figure 5A). Under glucose and glutamine replete conditions, the conversion of palmitate carbon into glutamate and aspartate was minimal (∼2-6%) (Figure 5D). However, a shift in palmitate carbon utilisation was observed in glucose and glutamine-deprived medium (∼10-19%) and to a lesser extent in glucose-deprived medium (∼5-14%) (Figure 3D). Overall, these results indicate that cancer cells can increase the catabolic conversion of palmitate into TCA cycle metabolites and amino acids, but only under extreme conditions when extracellular glucose or glucose plus glutamine are unavailable. Crucially, this shift in the labelling and distribution of fatty acid carbons represents a failed rescue.

While the fractional contribution of palmitate increased, the absolute abundance of TCA cycle intermediates (citrate, oxoglutarate, and malate) collapsed in glucose/glutamine-deprived conditions (Figure 5E, S3B). This indicates that even when reliance on FAO is forced by nutrient deprivation, fatty acids are inherently insufficient to replenish the TCA cycle. Importantly, these changes preceded the decrease in cell viability between 6 and 24 hours. The metabolic architecture we identified, in which FAO relies on glutamine anaplerosis (the Malic Enzyme shunt) to function, predicts this failure. Specifically, FAO-derived acetyl-CoA accumulates in the absence of an adequate anaplerotic substrate, stalling the TCA cycle and leading to metabolic collapse and cell death (Figure S5C). As such, these findings are consistent with our model, in which FAO can effectively contribute to TCA cycling only when glutamine-and/or glucose-derived anaplerosis is intact.

## Discussion

Cancer cells display remarkable metabolic plasticity, yet the quantitative contributions of individual fuels to mitochondrial metabolism have been difficult to establish in a comparative, pan-cancer context. While upregulation of FAO and sensitivity to CPT1 inhibition have been taken to imply that fatty acids are major mitochondrial fuels in many malignancies ^5,10,26,28–31^, yet direct measurements of fatty acid-derived carbon flux relative to glucose and glutamine have been lacking. By combining radiotracer assays, parallel stable-isotope tracing and quantitative mitochondrial phenotyping across 27 cancer cell lines, this study reveals that FAO capacity is profoundly dissociated from FA-derived TCA carbon flux, that FAO participates in a compensatory dual-fuel architecture with glutamine at the acetyl-CoA node, and that FAO cannot rescue TCA cycle activity or viability in the absence of glucose- and glutamine-derived anaplerosis. Understanding the specific contributions of fatty acids relative to glucose and glutamine in cancer metabolism could reveal novel vulnerabilities exploitable for targeted therapies.

Stoichiometrically, FAO generates significantly more ATP and reducing equivalents than other oxidative substrates ^26,27^. It is through this lens that much of the cancer FAO field has interpreted changes in gene expression and protein levels of those involved in FAO, as well as in vitro measures of fatty acid oxidative activity. For example, increased gene or protein levels of rate-limiting enzymes, or in vitro rates, are often interpreted as increased ATP-generating capacity and likely a dependency. Underpinning these interpretations is the assumption that an increase in capacity (i.e., in vitro flux) leads to greater carbon contribution to the TCA cycle and to more NADH and FADH_2_ to fuel the electron transport chain. Our results demonstrate a “capacity–contribution paradox” for FAO in cancer. Across a diverse panel of cell lines spanning 10 tissue origins, FAO rates varied by more than 8-fold, yet exogenous LCFAs contributed less than 10% of the carbon to TCA cycle intermediates. This pattern held across multiple FA species. By contrast, glucose and glutamine together accounted for the majority of TCA carbon, in line with their established roles as key mitochondrial fuels in cancer ^1–4,33^. Whilst basal and maximal FAO rates were related to CPT1A protein levels, cells with faster FAO rates had less mitochondrial mass. Further, spare FAO capacity in cells varied significantly across our panel, with cells that operated at higher basal FAO levels closer to their mitochondrial FAO ceiling and exhibiting limited relative plasticity to increase FAO flux.

In contrast, low basal FAO cells retain greater proportional reserve capacity despite lower absolute rates. These differences in FAO capacity and relationships with CPT1A protein levels and mitochondrial mass highlight the dangers in predicting FAO rates and mitochondrial fatty acid metabolic activity using these measures. Our findings necessitate a reinterpretation of prior studies that attributed the anti-tumour effects of CPT1 inhibition solely to blockade of a major fuel source^5,7–9,13–17^. There is strong clinical and preclinical interest in targeting FAO in a range of cancer types (see reviews^29–31^), and this remains well justified. However, the present data indicate that high FAO capacity does not imply that fatty acids supply most of the mitochondrial carbon. FAO cannot sustain TCA cycle activity or viability under conditions in which glucose- and glutamine-derived anaplerotic fluxes are acutely limited. Whilst previous studies have shown that FAO is upregulated when glycolysis is inhibited, or glucose is scarce ^24,34,35^, thereby suggesting that FAO provides a reserve fuel that can substitute for glucose under stress. Our data show increased FAO rates and fractional ¹³C palmitate labelling of TCA metabolites under glucose and glucose/glutamine deprivation, with absolute TCA metabolite pools collapsing and leading to reduced cell viability. These results demonstrate that FAO-derived acetyl-CoA can be used productively only when oxaloacetate supply from glucose and/or glutamine anaplerosis is sufficient. When both glucose and glutamine are limiting, FAO-derived acetyl-CoA cannot be efficiently condensed with oxaloacetate, leading to stalling of TCA cycling, depletion of intermediate pools, and increased diversion of fatty acid-derived carbons into cataplerotic products such as glutamate and aspartate. Therefore, FAO appears to support cancer metabolism through mechanisms beyond bulk carbon and ATP provision, including regulation of redox balance, fatty acyl-CoA pools for membrane homeostasis, or signalling lipids ^36–38^. Disentangling these non-canonical roles of FAO will be important for refining FAO-targeted therapies.

Having eliminated the carbon fuel role of FAO in cancer cells, we sought to reconcile why blocking fatty acid uptake ^19,20,23^ and metabolism ^22,39,40^ in mitochondria of cancer cells impacts cell viability. The inverse relationship between glucose- and FA-derived acetyl-CoA-CoA labelling recalls the classical Randle cycle described in muscle and adipose tissue ^32^, whereby high FAO suppresses PDH activity and thereby glucose oxidation. However, our data do not support this “FAO shuts down glucose oxidation” model. Firstly, glucose-derived anaplerosis via pyruvate carboxylase (m+3 citrate) is preserved across FAO states, indicating that glucose continues to refill TCA intermediates. Secondly, even in high-FAO cells, fatty acids remain minor contributors to TCA carbon, implying that any reduction in glucose-derived acetyl-CoA cannot be fully offset by FAO alone. Thirdly, a 5-fold increase in media fatty acid levels did not increase FAO rates; only lowering exogenous glucose levels to 3 mM or below increased FAO. As such, these observations argue for a reverse Randle Cycle^41^, whereby glucose availability and glycolytic flux dictate FAO rates.

We and many others have reported interactions between glucose and glutamine metabolism across diverse settings ^33,42–45^. These insights have showcased the critical importance of these nutrients in supporting cancer cell proliferation and spread. Here, we identified a compensatory “dual-fuel” architecture at the level of the mitochondrial acetyl-CoA pool, in which constraints on glucose-derived acetyl-CoA are jointly compensated by FAO and a glutamine-dependent malic enzyme shunt. Our data show that high-FAO cells display increased glutamine oxidation

(m+4 succinate) and enhanced labelling of downstream TCA intermediates and amino acids, consistent with elevated glutamine flux through canonical oxidative pathways. At the same time, glutamine-derived labelling of pyruvate and acetyl-CoA increases with FAO rate. Specifically, glutamine carbons enter the TCA cycle, pass through to malate, exit via malic enzyme to generate pyruvate, and re-enter as acetyl-CoA. This malic enzyme–dependent shunt aligns with prior observations that glutamine can be flexibly routed to support both anaplerosis and acetyl--CoA supply in various tumour contexts ^46–48^. Our results identify this as an attribute of cells with a high FAO metabolic state. This dual-fuel architecture provides an additional mechanistic explanation for the strong glutamine dependence observed in many cancers (see review^49^), yet may be more evident in cells with high FAO. As such, this new interpretation suggests that targeting glutamine metabolism is likely to be particularly effective in tumours with high FAO signatures; alternatively, in cells where FAO and glutamine metabolism cooperate to sustain acetyl-CoA when glucose-derived acetyl-CoA is limited.

Several limitations should be considered when interpreting our findings. First, the study focuses on exogenous long-chain fatty acids, glucose, and glutamine, which together account for 40–80% of TCA cycle labelling. Our results do not suggest that endogenous lipids and other nutrients, including branched-chain amino acids, are not important, as they will provide additional carbon^50–53^. Likewise, we used serum-free, FA-free BSA conditions to isolate the contributions of extracellular fatty acids; however, this approach does not fully recapitulate in vivo lipid availability, with its complex makeup of free fatty acids, lipoprotein-contained complex lipids, and those from local stromal sources ^50^. Finally, we acknowledge that all experiments were performed in 2D culture under controlled oxygen and nutrient conditions. In tumours, gradients of oxygen, pH, lipids and stromal interactions can modulate cancer cell metabolic activity. Currently, *in vivo* stable isotope tracing of fatty acid metabolism remains a major technical hurdle for the cancer metabolism field, as it requires significant circulating and intracellular precursor labelling that, to date, has been unachievable, especially to be stoichiometrically comparable to those achieved with ^13^C-glucose (>50% blood/plasma enrichment ^54–56^) and ^13^C-glutamine (30-60% blood enrichment ^55,57,58^).

In summary, this study provides quantitative evidence that exogenous long-chain fatty acids are minor carbon sources for the TCA cycle in cancer cells, irrespective of FAO capacity. Rather than acting as a primary mitochondrial fuel, FAO participates in a compensatory dual-fuel architecture in which fatty acid oxidation and a malic enzyme-dependent glutamine shunt together supplement acetyl-CoA synthesis when glucose-derived acetyl-CoA is low. The integrative nature of the metabolism of these nutrients is highlighted by the requirement for glucose to provide anaplerotic support, as FAO cannot maintain TCA cycle activity or cell viability in the absence of glucose- and glutamine-derived anaplerotic support. These findings collectively redefine the functional role of FAO in cancer metabolism, provide a mechanistic basis for reinterpretation of therapies targeting CPT1 and other FAO enzymes, and offer a framework for designing combination strategies that exploit the coupling between FAO, glutamine metabolism, and anaplerotic flux.

## Methods

### Key resources table

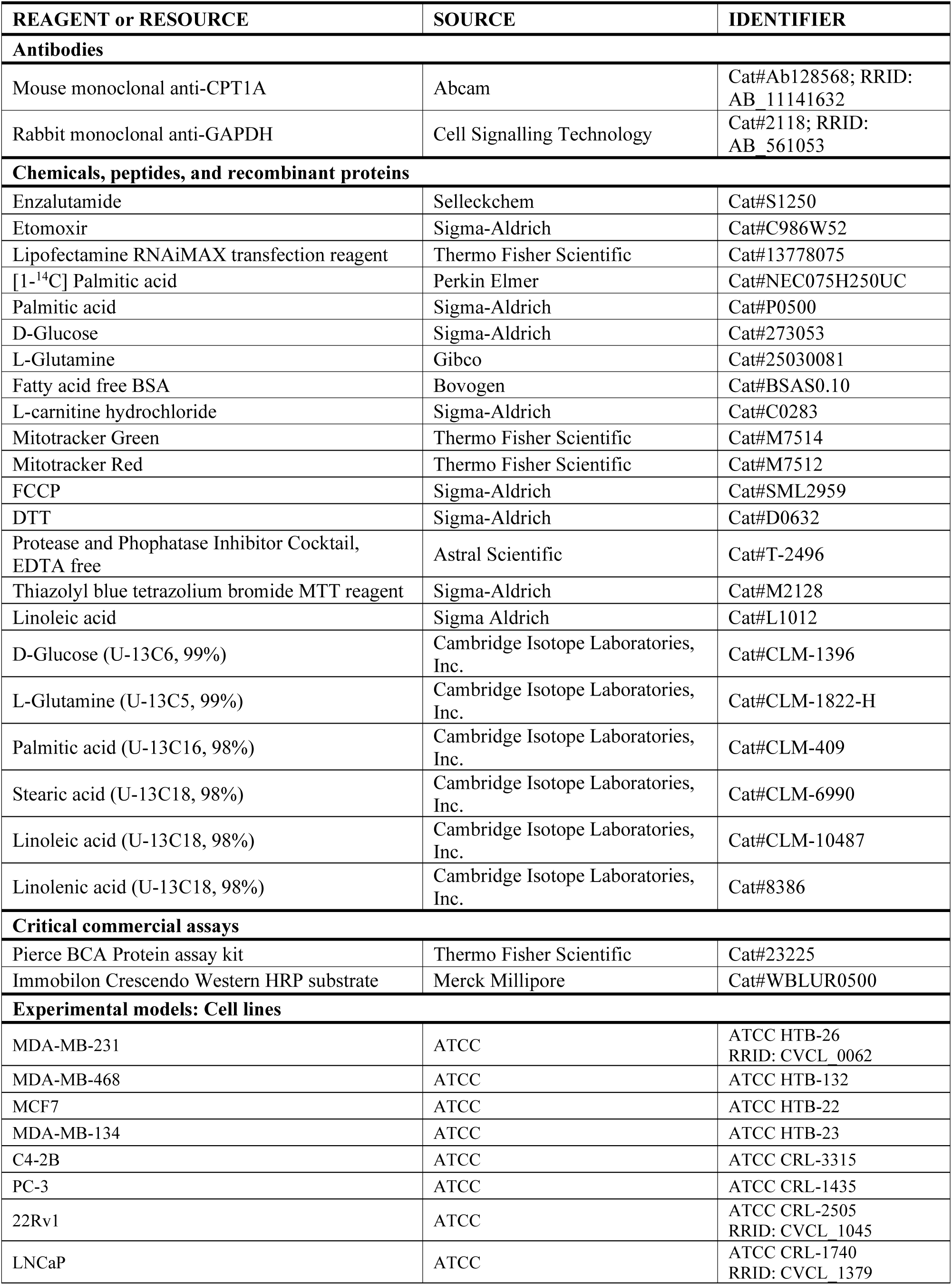
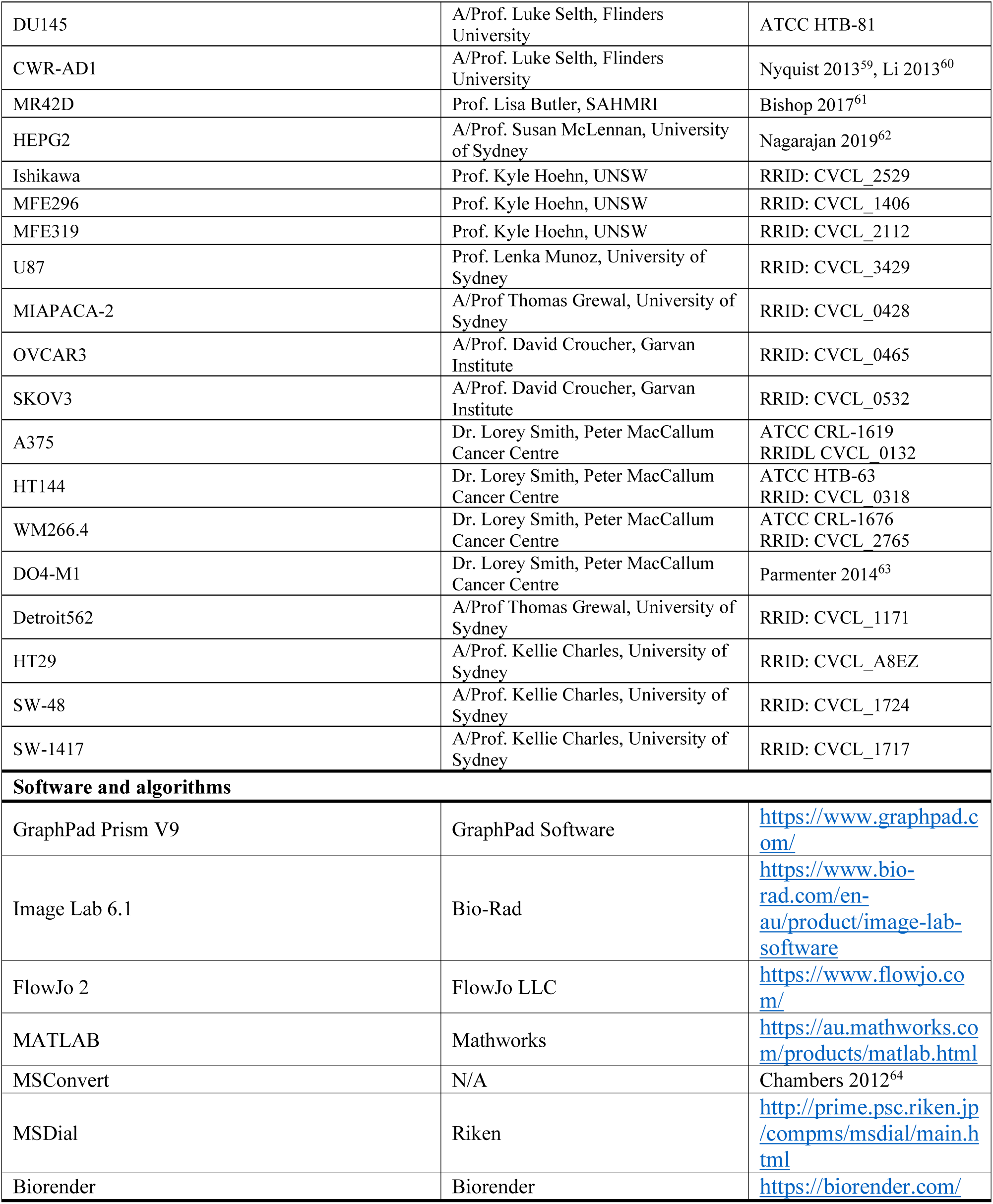

| REAGENT or RESOURCE | SOURCE | IDENTIFIER |
| --- | --- | --- |
| **Antibodies** |  |  |
| Mouse monoclonal anti-CPT1A | Abcam | Cat#Ab128568; RRID: AB 11141632 |
| Rabbit monoclonal anti-GAPDH | Cell Signalling Technology | Cat#2118; RRID: AB 561053 |
| **Chemicals, peptides, and recombinant proteins** |  |  |
| Enzalutamide | Selleckchem | Cat#S1250 |
| Etomoxir | Sigma-Aldrich | Cat#C986W52 |
| Lipofectamine RNAiMAX transfection reagent | Thermo Fisher Scientific | Cat#13778075 |
| [1-<sup>14</sup>C] Palmitic acid | Perkin Elmer | Cat#NEC075H250UC |
| Palmitic acid | Sigma-Aldrich | Cat#P0500 |
| D-Glucose | Sigma-Aldrich | Cat#273053 |
| L-Glutamine | Gibco | Cat#25030081 |
| Fatty acid free BSA | Bovogen | Cat#BSAS0.10 |
| L-carnitine hydrochloride | Sigma-Aldrich | Cat#C0283 |
| Mitotracker Green | Thermo Fisher Scientific | Cat#M7514 |
| Mitotracker Red | Thermo Fisher Scientific | Cat#M7512 |
| FCCP | Sigma-Aldrich | Cat#SML2959 |
| DTT | Sigma-Aldrich | Cat#D0632 |
| Protease and Phosphatase Inhibitor Cocktail, EDTA free | Astral Scientific | Cat#T-2496 |
| Thiazolyl blue tetrazolium bromide MTT reagent | Sigma-Aldrich | Cat#M2128 |
| Linoleic acid | Sigma Aldrich | Cat#L1012 |
| D-Glucose (U-13C6, 99%) | Cambridge Isotope Laboratories, Inc. | Cat#CLM-1396 |
| L-Glutamine (U-13C5, 99%) | Cambridge Isotope Laboratories, Inc. | Cat#CLM-1822-H |
| Palmitic acid (U-13C16, 98%) | Cambridge Isotope Laboratories, Inc. | Cat#CLM-409 |
| Stearic acid (U-13C18, 98%) | Cambridge Isotope Laboratories, Inc. | Cat#CLM-6990 |
| Linoleic acid (U-13C18, 98%) | Cambridge Isotope Laboratories, Inc. | Cat#CLM-10487 |
| Linolenic acid (U-13C18, 98%) | Cambridge Isotope Laboratories, Inc. | Cat#8386 |
| **Critical commercial assays** |  |  |
| Pierce BCA Protein assay kit | Thermo Fisher Scientific | Cat#23225 |
| Immobilon Crescendo Western HRP substrate | Merck Millipore | Cat#WBLUR0500 |
| **Experimental models: Cell lines** |  |  |
| MDA-MB-231 | ATCC | ATCC HTB-26<br>RRID: CVCL 0062 |
| MDA-MB-468 | ATCC | ATCC HTB-132 |
| MCF7 | ATCC | ATCC HTB-22 |
| MDA-MB-134 | ATCC | ATCC HTB-23 |
| C4-2B | ATCC | ATCC CRL-3315 |
| PC-3 | ATCC | ATCC CRL-1435 |
| 22Rv1 | ATCC | ATCC CRL-2505<br>RRID: CVCL 1045 |
| LNCaP | ATCC | ATCC CRL-1740<br>RRID: CVCL 1379 |
| DU145 | A/Prof. Luke Selth, Flinders University | ATCC HTB-81 |
| CWR-AD1 | A/Prof. Luke Selth, Flinders University | Nyquist 2013<sup>59</sup>, Li 2013<sup>60</sup> |
| MR42D | Prof. Lisa Butler, SAHMRI | Bishop 2017<sup>61</sup> |
| HEPG2 | A/Prof. Susan McLennan, University of Sydney | Nagarajan 2019<sup>62</sup> |
| Ishikawa | Prof. Kyle Hoehn, UNSW | RRID: CVCL_2529 |
| MFE296 | Prof. Kyle Hoehn, UNSW | RRID: CVCL_1406 |
| MFE319 | Prof. Kyle Hoehn, UNSW | RRID: CVCL_2112 |
| U87 | Prof. Lenka Munoz, University of Sydney | RRID: CVCL_3429 |
| MIAPACA-2 | A/Prof Thomas Grewal, University of Sydney | RRID: CVCL_0428 |
| OVCAR3 | A/Prof. David Croucher, Garvan Institute | RRID: CVCL_0465 |
| SKOV3 | A/Prof. David Croucher, Garvan Institute | RRID: CVCL_0532 |
| A375 | Dr. Lorey Smith, Peter MacCallum Cancer Centre | ATCC CRL-1619<br>RRID: CVCL_0132 |
| HT144 | Dr. Lorey Smith, Peter MacCallum Cancer Centre | ATCC HTB-63<br>RRID: CVCL_0318 |
| WM266.4 | Dr. Lorey Smith, Peter MacCallum Cancer Centre | ATCC CRL-1676<br>RRID: CVCL_2765 |
| DO4-M1 | Dr. Lorey Smith, Peter MacCallum Cancer Centre | Parmenter 2014<sup>63</sup> |
| Detroit562 | A/Prof Thomas Grewal, University of Sydney | RRID: CVCL_1171 |
| HT29 | A/Prof. Kellie Charles, University of Sydney | RRID: CVCL_A8EZ |
| SW-48 | A/Prof. Kellie Charles, University of Sydney | RRID: CVCL_1724 |
| SW-1417 | A/Prof. Kellie Charles, University of Sydney | RRID: CVCL_1717 |
| **Software and algorithms** |  |  |
| GraphPad Prism V9 | GraphPad Software | https://www.graphpad.com/ |
| Image Lab 6.1 | Bio-Rad | https://www.bio-rad.com/en-au/product/image-lab-software |
| FlowJo 2 | FlowJo LLC | https://www.flowjo.com/ |
| MATLAB | Mathworks | https://au.mathworks.com/products/matlab.html |
| MSConvert | N/A | Chambers 2012<sup>64</sup> |
| MSDial | Riken | http://prime.psc.riken.jp/compms/msdial/main.html |
| Biorender | Biorender | https://biorender.com/ |

### Method details

#### Cell lines and culture conditions

A panel of 27 adherent cancer cell lines of 10 different origins was used in this study. Cell lines were obtained from both vendors and kindly provided by academic laboratories. Details of culture media (Gibco) can be found in Supplementary Table 1. All culture media were supplemented with 10% fetal bovine serum (Cytiva Hyclone) and 1% penicillin/streptomycin (Gibco). Cells were incubated at 37°C and 5% CO_2_.

#### 14C-Radiolabelling

Cells were seeded in technical triplicate in 6-well plates. The next day, the cells were washed in warm PBS and incubated in 600 μL media containing 0.1 mmol/L cold palmitate, [1-^14^C] palmitate (0.1 μCi/mL; Perkin Elmer) conjugated to 2% (wt/vol) FA-free BSA and 1 mmol/L -carnitine in RPMI or DMEM glucose-free medium (Gibco) for 4 hours. Fatty acid oxidation rates were determined from CO_2_ production as previously described^19^ in either 0 mM or 5 mM glucose to measure FAO flexibility. Protein content was measured by BCA protein assay and absorbance was read at 540 nm using a Tecan infinite M1000 pro plate reader.

#### Radiolabelled glucose titration

MDA-MB-231 cells were treated as described above for radiolabelled experiments, and supplemented with 0, 1, 3, 5, 7, 9 11, and 25 mM glucose +/- 10 μM etomoxir. In the selected panel of six cell lines, cells were treated as described above for radiolabelled experiments, supplemented with 0, 1, 3, and 5 mM glucose.

#### Mitotracker

Cells were seeded in 12-well plates at a density of 2×10^5^ cells/well in RPMI or HG DMEM medium. After 24 hours, cells were washed in warm PBS, trypsinised, and collected in serum-free RPMI medium. The cell suspension was centrifuged at 1500 rpm for 5min in fluorescence-activated cell sorting (FACS) tubes. Cells were subsequently stained with 200 nM Mitotracker Red and Mitotracker Green at 37°C for 30 mins. Additionally, unstained, red only, green only, and 2.5mM FCCP treated cells were included as experimental controls. Cells were centrifuged at 1500 rpm for 5min and fixed with 4% paraformaldehyde at 37°C for 10 mins. Cells were washed with warm PBS, centrifuged at 1500 rpm for 5 mins and resuspended in 150 mL PBS before the samples were read on a BD LSR Fortessa flow cytometer. Data analysis was performed using FlowJo software.

#### Immunoblotting

Cells were seeded in 6-well plates (5×10^5^). After 24 hours, cells were lysed in RIPA buffer containing 1% DTT and 0.1% protease and phosphatase inhibitor cocktail. Protein content was measured by BCA protein assay, and absorbance was read at 540 nm using a TECAN infinite M1000 pro plate reader. Cell lysates were subjected to SDS-PAGE, and transferred to polyvinylidene difluoride (PVDF) membranes (Merck Millipore). Ponceau staining was performed on the membranes as a loading control, and then membranes were immunoblotted for CPT1a. Chemiluminescence was performed using Luminata Crescendo Western HRP Substrate and imaged using the Bio-Rad ChemiDoc MP Imaging System and Image Lab software 4.1 (Bio-Rad Laboratories).

#### Cell viability

Cells were seeded in 96-well plates in 6 well replicates (3×10^3^ cells/well). After 24 hours the media was removed and replaced with glucose- and glutamine free media (Gibco) supplemented with 5 mM glucose, 1 mM glutamine and/or 150 µM fatty acids, conjugated to 2% (wt/vol) FA-free BSA. For glucose-deprived conditions, DMEM no glucose, no glutamine media was supplemented with 1 mM glutamine and 150 μM palmitate; for glucose and glutamine-deprived conditions, media was supplemented with 150 µM palmitate only. Media supplemented with 10% FBS was used as a control. At 6, 12, and 24 hours, 50 µl of MTT reagent was added to each well. MTT reactions were incubated at 37°C and 5% CO_2_ for 4 hours, and then stopped by replacing the media with 100 µl DMSO (Sigma–Aldrich) per well. Absorbance was read at 540 nm using a TECAN infinite M1000 Pro plate reader. Cells were seeded in 6-well plates at 5×10^5^ cells/well for live cell count. After 24 hours, media was replaced as described for the MTT assay, and the cells counted by trypan blue (Gibco) exclusion at 6 and 24 hours.

#### [U-^13^C]-stable isotope tracing

Cells were seeded in 6-well plates (5×10^5^). After 24 hours, plates were washed with warm PBS and media replaced with 600 μL of no glucose and no glutamine DMEM (Gibco) supplemented with 5 mM glucose, 1 mM glutamine and/or 150 μM FAs, replaced with their respective [U-^13^]-C forms. [U-^13^C]-fatty acids used were palmitate (16:0), oleate (18:0), stearate (18:1), linoleate (18:2), and linolenate (18:3). For glucose-deprived conditions, no glucose and no glutamine DMEM was supplemented with 1 mM glutamine and 150 μM [U-^13^C]-palmitate; for glucose and glutamine-deprived conditions media was supplemented with 150 μM [U-^13^C]-palmitate only. Fatty acids were conjugated to 2% (wt/vol) fatty acid-free BSA-containing media for 24 hours at 37 °C before treating cells. Cells were incubated in ^13^C containing medium for 6 hours.

#### Extracellular substrate sampling

Cells were seeded in 6-well plates (5×10^5^). After 24 hours, the wells were washed with warm PBS and media replaced with 1 mL or 2 mL of DMEM with no glucose, no glutamine media supplemented with 5 mM glucose, 1 mM glutamine, and 150 μM fatty acids, conjugated to 2% (wt/vol) FA-free BSA. 100 μL of extracellular media was collected from the wells at 3, 6, 12, and 24 hour timepoints for cells incubated in 1 mL of media, and 12 and 24 hours for cells incubated in 2 mL of media. Media samples were centrifuged at 16,000 x g for 5 mins at 4°C, and the supernatant was collected for subsequent LC-MS analysis.

#### Cell lysate extraction for LC-MS

Media was aspirated from the plates and the cells washed with 2 mL ice-cold 0.9% (w/v) NaCl. Cells were scraped with 300 μL of extraction buffer, EB, (1:1 LC/MS methanol:water (Optima) + 0.1x internal standards (non-endogenous polar metabolites) and transferred to a 1.5 mL microcentrifuge tube. A further 300 μl of EB was added to the cells and combined in the tube. Next, 600 μL of chloroform (Honeywell) was added before vortexing and incubating on ice for 10 mins. Tubes were vortexed briefly and centrifuged at 15,000g for 10 mins at 4°C. The aqueous upper layer was collected and dried without heat using a Savant SpeedVac (Thermo Fisher) for metabolomics LC-MS analysis. The lower layer was collected and dried under nitrogen flow for lipidomic LC-MS analysis.

#### Measuring stable-isotope labelled metabolites by LC-MS

Dried aqueous upper phase samples were resuspended in 40 mL amide buffer A (20mM ammonium acetate, 20 mM ammonium hydroxide, 95:5 HPLC H_2_O: Acetonitrile (v/v)) and vortexed and centrifuged at 15 000 x g for 5 mins at 4°C. 20 mL of supernatant was transferred to HPLC vials containing 20 mL acetonitrile for LCMS analysis of amino acids and glutamine metabolites. The remaining 20 mL of the resuspended sample was transferred to HPLC vials containing 20 mL of LC-MS H_2_O for LCMS analysis of glycolytic, pentose phosphate pathway, and TCA cycle metabolites. Amino acids and glutamine metabolites were measured using a Vanquish-TSQ Altis (Thermo) LC-MS/MS system. Analyte separation was achieved using a Poroshell 120 HILIC-Z Column (2.1×150 mm, 2.7 mm; Agilent) at ambient temperature. The pair of buffers used were amide buffer A and 100% acetonitrile (Buffer B), flowed at 200 mL/min; injection volume of 5 mL. Glycolytic, pentose phosphate pathway and TCA cycle metabolites were measured using 1260 Infinity (Agilent)-QTRAP6500+ (AB Sciex) LC-MS/MS system. Analyte separation was achieved using a Synergi 2.5 mm Hydro-RP 100A LC Column (100×2 mm) at ambient temperature. The pair of buffers used were 95:5 (v/v) water:acetonitrile containing 10 mM tributylamine and 15 mM acetic acid (Buffer A) and 100% acetonitrile (Buffer B), flowing at 200 mL/min; injection volume of 5 mL. Raw data from both LC-MS/MS systems were extracted using MSConvert^64^ and in-house MATLAB scripts.

#### Measuring extracellular substrates by LC-MS

20 μL of collected extracellular media was mixed with 80 μL water, and vortexed. 10 μL of diluted media was mixed with 90 μL of extraction buffer containing 1:1 (v/v) acetonitrile and methanol + 1x internal standards (non-endogenous standards) at −30°C. The mixture was centrifuged at 12,000xg for 5 min at 4°C and transferred into HPLC vials for LC-MS analysis measured using the Vanquish-TSQ Altis as described above. Raw data from both LC-MS/MS systems were extracted using MSConvert^64^ and in-house MATLAB scripts.

#### Measuring stable isotope labelled lipids by LC-MS

Dried lower phase samples were resuspended in 100 μL 4:2:1 (v/v) isopropanol:methanol:chloroform containing 7.5 mM ammonium formate. Acyl-carnitine lipids were measured using a Thermo Scientific Q-Exactive-HF-X Hybrid Quadrupole Orbitrap LC-MS/MS system. Analyte separation was achieved using an Agilent Poroshell 120, HILIC, 2.1×100 mm, 2.7 μm column. The pair of buffers used were 0.1(v/v)% formic acid and 10 mM ammonium formate in H_2_O and 0.1(v/v)% formic acid in acetonitrile, flowed at 200 μL/min on positive mode. MS1 data was acquired with the following settings: 3.5 kV, capillarity temperature 300°C, 120,000, injection time 100 ms, AGC 1×10^6^, scan range 200-1000. For ddMS2 data acquisition, the following settings were used: top 5, resolution 30,000, 200-2000, isolation 1.0m/z, nce 30, AGC target 1×10^3^, intensity threshold 5×10^3^, dynamic exclusion 20s. Raw data from both LC-MS/MS systems were extracted using MSConvert^64^ and in-house MATLAB scripts.

#### Quantification and statistical analysis

Student’s t-test, one- or two-way ANOVA with Dunnett’s or Tukey’s multiple comparisons, and linear regression statistical tests were performed using GraphPad Prism 11.0.0 (GraphPad Software). *P* < 0.05 was considered significant. Data are reported as mean ± standard error of the mean (SEM), standard deviation (SD), or min-max values of at least three independent determinations. Radiolabelling experiments for the expansive cell line panel were of at least one independent determination (N=1-x as indicated in the respective figure legend) with at least 3 technical replicates each. Schematic diagrams were created using Biorender.com software.

## Supplementary information

**Supplementary Table 1:**
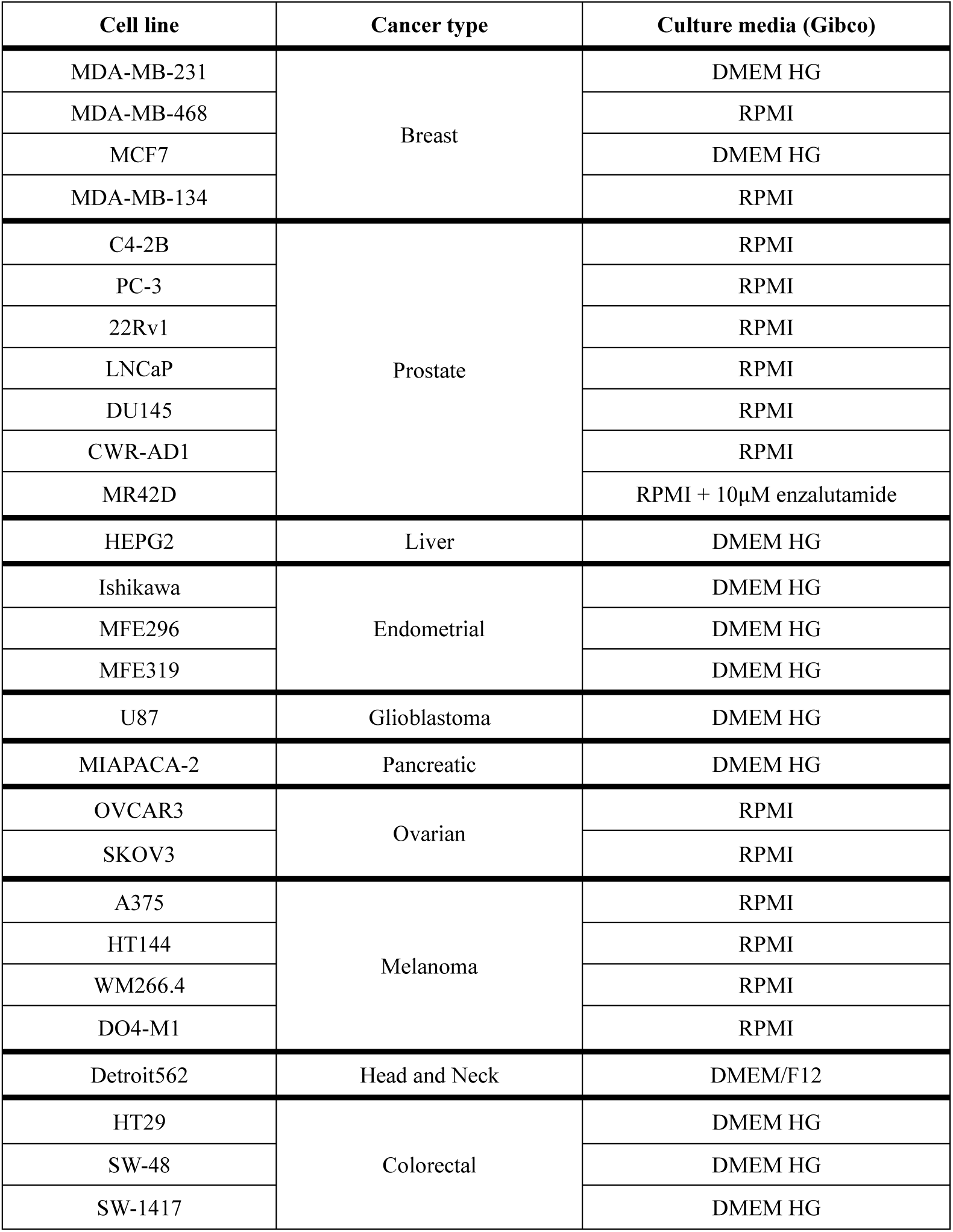
Cell line details.

| Cell line | Cancer type | Culture media (Gibco) |
| --- | --- | --- |
| MDA-MB-231 | Breast | DMEM HG |
| MDA-MB-468 |  | RPMI |
| MCF7 |  | DMEM HG |
| MDA-MB-134 |  | RPMI |
| C4-2B | Prostate | RPMI |
| PC-3 |  | RPMI |
| 22Rv1 |  | RPMI |
| LNCaP |  | RPMI |
| DU145 |  | RPMI |
| CWR-AD1 |  | RPMI |
| MR42D |  | RPMI + 10µM enzalutamide |
| HEPG2 | Liver | DMEM HG |
| Ishikawa | Endometrial | DMEM HG |
| MFE296 |  | DMEM HG |
| MFE319 |  | DMEM HG |
| U87 | Glioblastoma | DMEM HG |
| MIAPACA-2 | Pancreatic | DMEM HG |
| OVCAR3 | Ovarian | RPMI |
| SKOV3 |  | RPMI |
| A375 | Melanoma | RPMI |
| HT144 |  | RPMI |
| WM266.4 |  | RPMI |
| DO4-M1 |  | RPMI |
| Detroit562 | Head and Neck | DMEM/F12 |
| HT29 | Colorectal | DMEM HG |
| SW-48 |  | DMEM HG |
| SW-1417 |  | DMEM HG |

## Acknowledgements

N.T.S. was supported by the Australian Rotary Health/Rotary Club of Blacktown City ‘Mel Grey’ PhD scholarship. A.J.H. was supported by a Robinson Fellowship from the University of Sydney. This study was supported by funding from The University of Sydney. We thank the Sydney Mass Spectrometry facility for access to LC-MS instruments; Associate Professor Helen McGuire and the Sydney Cytometry core facility for assistance with MitoTracker experiments; and all collaborators who kindly donated cell lines specified in the Key Resources Table.

## Author contributions

Conceptualisation by A.J.H. and L.E.Q. Project administration, visualisation, writing-original draft by N.T.S. Investigation by N.T.S., M.F.H.S., M.G., and A.W. Methodology by L.E.Q. Software by N.T.S., and L.E.Q. Formal analysis by N.T.S, L.E.Q., and A.J.H. Writing-editing & reviewing by N.T.S., L.E.Q., and A.J.H. Funding acquisition, resources by A.J.H. Supervision by A.J.H., and L.E.Q.

## Availability of data and materials

All data generated or analysed during this study are included in this published article and its supplementary information files. The datasets generated during the current study are available from the corresponding author upon reasonable request.

